# A framework for ultra-low input spatial tissue proteomics

**DOI:** 10.1101/2023.05.13.540426

**Authors:** Anuar Makhmut, Di Qin, Sonja Fritzsche, Jose Nimo, Janett König, Fabian Coscia

## Abstract

Spatial tissue proteomics combining microscopy-based cell phenotyping with ultra-sensitive mass spectrometry (MS)-based proteomics is an emerging and powerful concept for the study of cell function and heterogeneity in health and disease. However, optimized workflows that preserve morphological information for image-based phenotype discovery and maximize proteome coverage of few or even single cells from laser microdissected archival tissue, are currently lacking. Here, we report a robust and scalable workflow for the proteomic analysis of ultra-low input formalin-fixed, paraffin-embedded (FFPE) material. Benchmarking in the murine liver resulted in up to 2,000 quantified proteins from single hepatocyte contours and nearly 5,000 proteins from 50-cell regions with high quantitative reproducibility. Applied to human tonsil, we profiled 146 microregions including spatially defined T and B lymphocyte niches and quantified cell type specific markers, cytokines, immune cell regulators and transcription factors. These rich data also highlighted proteome dynamics in spatially defined zones of activated germinal centers, illuminating sites undergoing active B-cell proliferation and somatic hypermutation. Our results demonstrate the power of spatially-resolved proteomics for tissue phenotyping by integrating high-content imaging, laser microdissection, and ultra-sensitive mass spectrometry. This approach has broad implications for a wide range of biomedical applications, including early disease profiling, drug target discovery and biomarker research.

## INTRODUCTION

Cells are the functional units of organs, which fulfill essential physiological tasks in a spatially defined manner to maintain tissue integrity ^1^. To analyze cell dynamics in space and time, powerful spatial genomics ^2^, epigenomics ^3^, transcriptomics ^4–6^ and imaging-based proteomics ^7, 8^ methods have been developed to better understand cellular and molecular drivers of health and disease states. As proteins are the biomolecules closest to the cellular phenotype determining cell identity and function ^9, 10^, spatial proteomics (SP) methods are particularly promising for the study of human (patho)physiology. SP methods with the single-cell resolution are dominated by targeted antibody-based methods such as imaging mass cytometry ^11^ (IMC) or multiplex immunofluorescence (mIF) imaging ^8, 12^, where several dozen proteins can be analyzed at (sub)cellular resolution. However, while such methods are well-suited for the large-scale screening of cellular phenotypes, they fall far short in capturing the actual complexity of the cellular proteome. It is estimated that single cell types express more than 10,000 unique proteins ^9^, which is complemented by millions of potential proteoforms, including splice variants, post-translational modifications (PTMs), and protein sequence variants ^10, 13, 14^. Liquid chromatography mass spectrometry (LC-MS) based proteomics in contrast enables the study of proteomes at an unbiased (i.e., untargeted), quantitative and system-wide level ^9^. The combination of both of these complementary proteomic approaches is therefore highly desirable, but requires integrated and multimodal pipelines. We recently introduced Deep Visual Proteomics (DVP)^15^, a new concept combining imaging-based (fluorescence or brightfield) single-cell phenotyping with unbiased MS-based proteomics for global proteome profiling with cell type and spatial resolution. To realize DVP, we developed an automated laser microdissection (LMD) workflow for the streamlined collection of nuclei, cells or larger regions of interest (ROI) directly into 96 or 384-well plates, thereby connecting whole-slide imaging and deep-learning-based image analysis ^16^ with ultra-sensitive MS-based proteomics ^17^. This allowed the profiling of as little as 100 phenotype-matched cells from archival tissue material, while also preserving detailed cell type and spatial information. Further advances in sample preparation and MS acquisition recently pioneered the profiling of single-cell proteome heterogeneity in cryosections of murine liver tissue ^18^, emphasizing the strong spatial influence on the hepatocyte-specific proteome. Despite these promising proof-of-concept studies, a systematic evaluation and optimization of all experimental steps of immunofluorescence microscopy-guided spatial tissue proteomics is still missing. In particular, the analysis of few or even single cells of FFPE tissue collected by laser microdissection has remained elusive and relies on optimized and robust ‘end-to-end’ protocols. The successful development of such integrated workflows could pave the way for a plethora of biomedical applications, including early disease proteome profiling studies directly from archived patient material, where only a few cells can be present.

Here, we describe a scalable, robust and easy-to-use protocol optimized for the profiling of ultra-low input archival tissue guided by whole-slide (immunofluorescence) imaging. After benchmarking in murine liver tissue, we applied our workflow to study the cell type resolved proteome of B and T lymphocytes in different spatially defined niches, guided by four-marker whole-slide immunofluorescence imaging. We finally provide detailed guidelines covering all aspects of our workflow, from antigen retrieval and staining on LMD membrane slides, over manual or automated sample processing, to MS data acquisition and analysis.

## RESULTS

### Optimizing laser microdissection-based low-input FFPE tissue proteomics

Light microscopy is an integral part of spatial proteomics workflows that combine LMD with MS-based proteomics ^19–21^ (**Fig. 1a**). Whereas hematoxylin and eosin (H&E) staining protocols are well-established for tissue sections mounted on specialized LMD membrane slides ^22^ (**Extended Data Fig. 1a**), immunofluorescence-based protocols, which allow the sensitive detection of a higher number of cell type and functional markers (typically 3-5 per image cycle), require additional antigen retrieval (AR) steps for epitope unmasking. The choice of the AR method not only influences staining and image quality, but also impacts LMD membrane integrity, tissue collection efficiency and potentially proteome coverage of trace sample amounts. The evaluation and optimization of AR and staining prior to MS is therefore important for ultra-low input or even single-cell applications that strictly depend on highly efficient tissue collection and protein extraction, as well as near-lossless sample preparation protocols. In other words, the most sensitive mass spectrometer can only be as good as the upstream sample preparation workflow that delivers these trace peptide amounts to the instrument. We therefore first evaluated different AR methods for their general compatibility with whole-slide imaging on LMD slides, efficient LMD and ultra-sensitive MS-based proteomics. We tested two common protocols based on heat-induced (HIER) or proteolytic (i.e., Pepsin-induced, PIER) epitope retrieval and immunofluorescently stained murine liver FFPE tissue with an antibody against an ubiquitously expressed plasma membrane marker (Na/K ATPase). We chose murine liver as benchmarking tissue as hepatocytes make up 60-80% of the liver cell mass ^23^, which allowed us to obtain consistent results from serial, homogenous tissue slices and repeated sample collections over the course of our experiments. 5 µm-thick liver sections were mounted on glass membrane (PEN or PPS) or metal frame (PPS) slides, de-paraffinized and subjected to two common antigen retrieval protocols (Methods) prior to antibody staining, immunofluorescence microscopy, LMD and proteomics (**Fig. 1b**). We included H&E stains for comparison, which do not rely on additional antigen retrieval after de-paraffinization, allowing us to directly investigate the impact of PIER and HIER on our proteomic results. PIER was fully compatible with both LMD slide types (glass or metal frame), facilitating efficient laser microdissection, but generally came at the cost of higher background staining compared to HIER (**Fig. 1c**). For macroscopic sample acquisition with glass slides HIER on the contrary occasionally resulted in membrane distortion of the glass type slides towards the label end of the slide (**Extended Data Fig. 1a**), which can impede efficient LMD collection in this area if not otherwise prevented ^24^. Frame slides are applicable to many biological sample types (i.e., tissue or cell culture), but are suboptimal for the collection of directly connected contours, for example for gridded sampling schemes as used for the nanoPOTS approach ^20^, as this can ultimately lead to loss of overall membrane integrity. Importantly, irrespective of the choice of antigen retrieval (HIER or PIER), staining technique (H&E or IF), or LMD slide type (PEN or PPS), proteomics results from three different liver tissue amounts were highly consistent. Tissue samples were processed in 384-well low-binding plates using an MS-compatible, organic solvent-based protocol, which included 60-min heating at 95°C for efficient formalin decrosslinking ^25^, sequential Lys-C and trypsin digestion and miniaturized solid-phase extraction. Using 2 µl lysis buffer, our workflow was easily pipette-able with standard laboratory equipment, but at the same time also allowed the integration of robotic sample preparation workflows. Using an optimized 15-min active nano-LC gradient (**Extended Data Fig. S1b-d**) in combination with dia-PASEF ^26^ on a trapped-ion mobility spectrometry (TIMS) mass spectrometer (Bruker timsTOF SCP) and DIA-NN ^27^, we quantified more than 9,000 precursors and 1,700 unique proteins from small tissue regions of approximately one to two hepatocytes (∼1,500 µm^2^, 5 µm thick, **Fig. 1d-e, Extended Data Fig. 1e-f**). From 50-cell samples (50,000 µm^2^, 5µm thick) of HE, HIER and PIER-derived samples, 4,000 proteins were consistently quantified with excellent quantitative reproducibility (Pearson r = 0.98, **Fig. 1d**). Furthermore, we found a 95% overlap of protein identifications across AR methods (**Fig. 1f**), as well as nearly identical cellular compartment proportions of the underlying proteomes (**Fig. 1g**). The comparison of our FFPE data to a deep liver proteome study in primary cells ^28^, also showed similar cellular compartment distributions. We conclude that common FFPE tissue preparation methods for fluorescence microscopy are fully compatible with ultra-low input MS-based proteomics. Based on our data included here and input from two additional LMD expert labs, we further summarized these findings in **Fig. 1h** to provide general guidelines for LMD proteomics beginners. Together with our detailed protocol description (**Methods and Supplementary information**), these could support the selection of the right tissue preparation and sample collection strategy for diverse spatial proteomics applications.

**Fig. 1:**
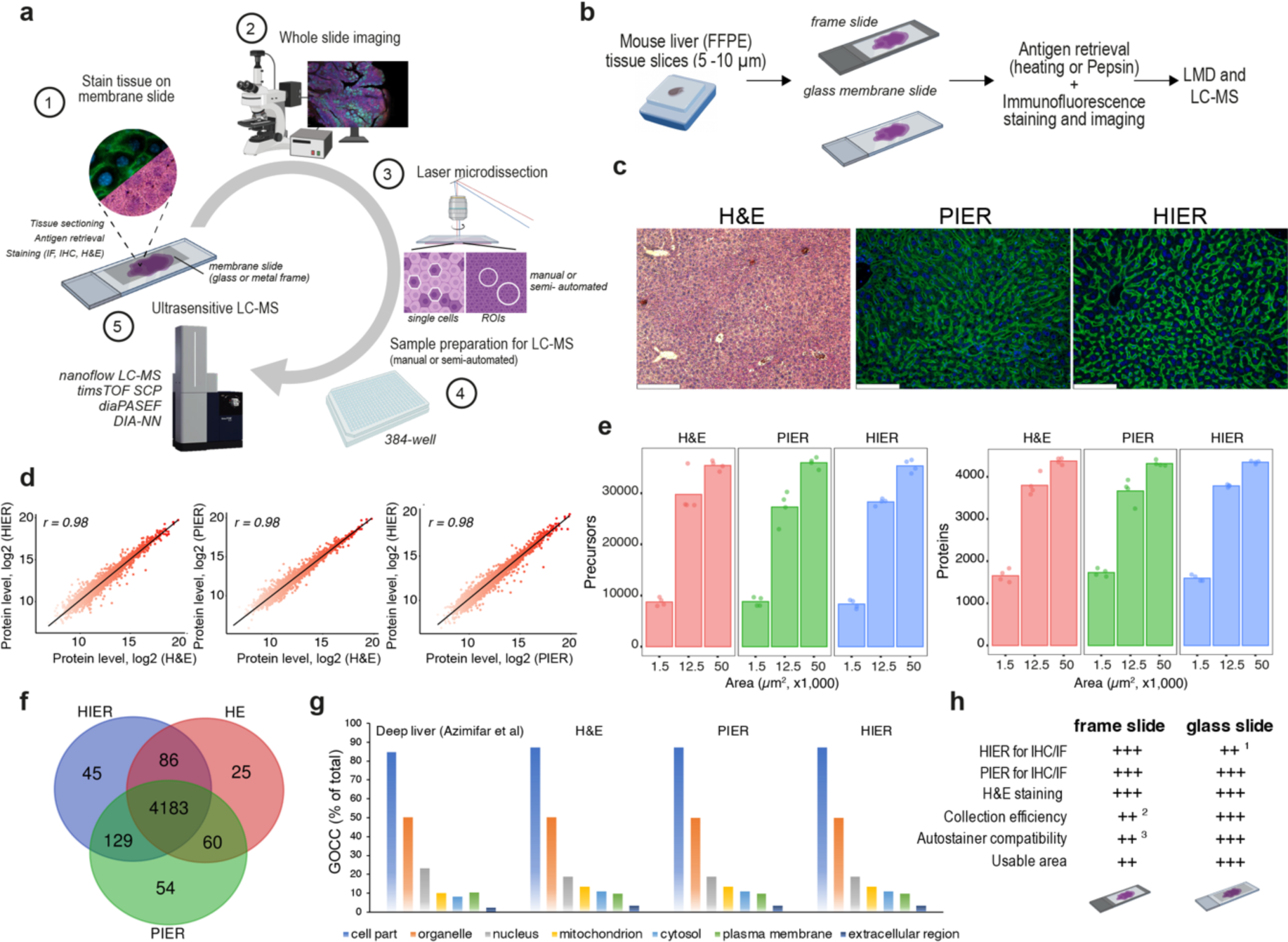
Guiding laser microdissection-based tissue proteomics. **a)** Overview of the spatial tissue proteomics workflow. **b)** Tissue preparation strategy for laser microdissection-based proteomics benchmarking experiments using specialized LMD slides (metal frame or glass). **c)** Representative images of hematoxylin and eosin (H&E) or immunofluorescence-stained mouse liver based on different antigen retrieval methods. PIER: pepsin-induced epitope retrieval, HIER: heat-induced epitope retrieval. Scale bars: Left: 249 µm, middle and right 124.5 µm. **d)** Proteome correlations of HIER, PIER and HE-based tissue samples. Pearson correlations. **e)** Precursor and protein identifications from tissue samples processed with different staining and antigen retrieval methods. Areas of 1,500, 12,500 and 50,000 µm^2^ of a 5-µm thick section were laser microdissected. Averages are shown from quadruplicate measurements. **f)** Venn diagram showing common and exclusive proteins for different antigen retrieval strategies based on 50,000 µm^2^ tissue samples. **g)** Protein identifications from major cellular compartments (‘cytosol’, ‘nucleus’, ‘plasma membrane’ and ‘extracellular region’, Gene Ontology Cellular Component, GOCC) from HE, PIER and HIER-treated mouse liver tissues. Percentages are the number of quantified proteins per compartment over all quantified proteins in the corresponding sample. For comparison, a deep mouse liver proteome dataset was included ^28^ based on non-fixed, primary cells. **h)** Summary of the laser-microdissection optimizations for low-input proteomics. Three labs assessed the applicability of glass and frame slides and rated each category with moderate (+), good (++) and excellent (+++). The average score is shown from all three ratings. ^1^HIER can cause membrane distortion of glass-type slides. ^2^Frame slides are more problematic for grid-based sampling schemes, the collection of many closely connected contours can lead to loss of overall membrane integrity. ^3^Tested autostainers: Ventana (Roche) supported frame and glass slides, DAKO (Agilent) system supported glass slides. Panels a and b created with Biorender.

### A scalable FFPE tissue proteomics workflow allowing single-cell analysis

Having established that common antigen retrieval and staining methods are fully compatible with laser microdissection and ultra-low input tissue proteomics, we next assessed the scalability, robustness and minimally required sample amount of our workflow. Homogenous areas of murine liver tissue were collected by laser microdissection into 384-well low-binding plates, ranging from single hepatocytes (600 µm^2^, 5 µm thick) to approx. 100 cells (100,000 µm^2^, **Fig. 2a-b**, **Extended Data Fig. 1e-f**). Proteomic results showed a linear increase in MS2 intensity, precursor and protein identifications from ‘low’ to ‘high’ tissue amounts, which, as expected, was inversely correlated with the median coefficient of variations (CV) of protein quantifications calculated from triplicate measurements (**Fig. 2c-e**). However, even single hepatocyte contours featured low median CVs of 20% for the 1,500 – 2,000 quantified proteins per contour (**Fig. 2e, Extended Data Fig. 2a-b**), which we and others previously only achieved from many thousands of cells collected from FFPE tissue ^29, 30^. It is noteworthy that CVs likely also included true biological variation from known spatially defined hepatocyte heterogeneity ^18, 31^, letting us conclude that reproducible single-cell proteome analysis from FFPE tissue is readily achievable based on this workflow. We attribute the excellent quantitative reproducibility to the combination of optimized tissue preparation, low-volume sample processing and highest-sensitivity MS acquisition using an optimal window dia-PASEF scheme (Methods). Precursor and protein identifications peaked for 50-100 cell regions (25,000 – 50,000 µm^2^), where we quantified close to 50,000 precursors and 5,000 proteins (**Extended Data Table 1**). Interestingly, further increasing sample amounts resulted in lower precursor identifications and higher median CVs, indicative of sample overloading (**Fig. 2d-e**). While this phenomenon could possibly be balanced out using longer LC gradients, thereby further improving proteome coverage, we note that our optimized 15-min active nanoflow gradient provides an excellent compromise between single-cell sensitivity, high proteome coverage for 50-100 cell samples, and reasonable sample throughput of around 30-40 samples per day. Additionally, we tested the applicability of our protocol in combination with the Bruker timsTOF Pro2, which shows a roughly four to five-fold lower total ion current compared to the SCP instrument ^32^. For the lowest tissue amounts measured (7,500 µm^3^ samples, 1-2 hepatocytes ^33^), close to 1,000 proteins could still be quantified with high quantitative reproducibility and a somewhat similar proteome coverage for the 50-cell samples (**Extended Data Fig. 2c-e**). We next tested if we could further increase proteome coverage of single FFPE tissue contours through the use of MS-compatible detergents. We assessed the integration of n-Dodecyl-β-D-maltoside (DDM), which was previously shown to improve proteome coverage of low-input FFPE tissue, in particular when applied at high temperatures ^34^. Combining our acetonitrile (ACN) based protocol (10% final conc.) with DDM (0.1% final conc.), which likewise included 60-min controlled heating at 95°C for efficient formalin de-crosslinking, proteome coverage of the single-cell samples improved by 104% and 50% for precursor and protein identifications, respectively (**Extended Data Fig. 2g**). For 12,500 µm^2^ (∼12 cells) or 50,000 µm^2^ (∼50 cells) samples, proteome coverage was more similar between protocols. As DDM is not removed during peptide clean-up steps, its accumulation on the analytical column can compromise chromatographic performance over time, which can be prevented by additional high organic solvent washing. Our organic solvent-based protocol performed equally well for 50-cell contours, thus offering an excellent and cleaner alternative to the DDM/ACN combination for ‘higher’ tissue amounts.

**Figure 2:**
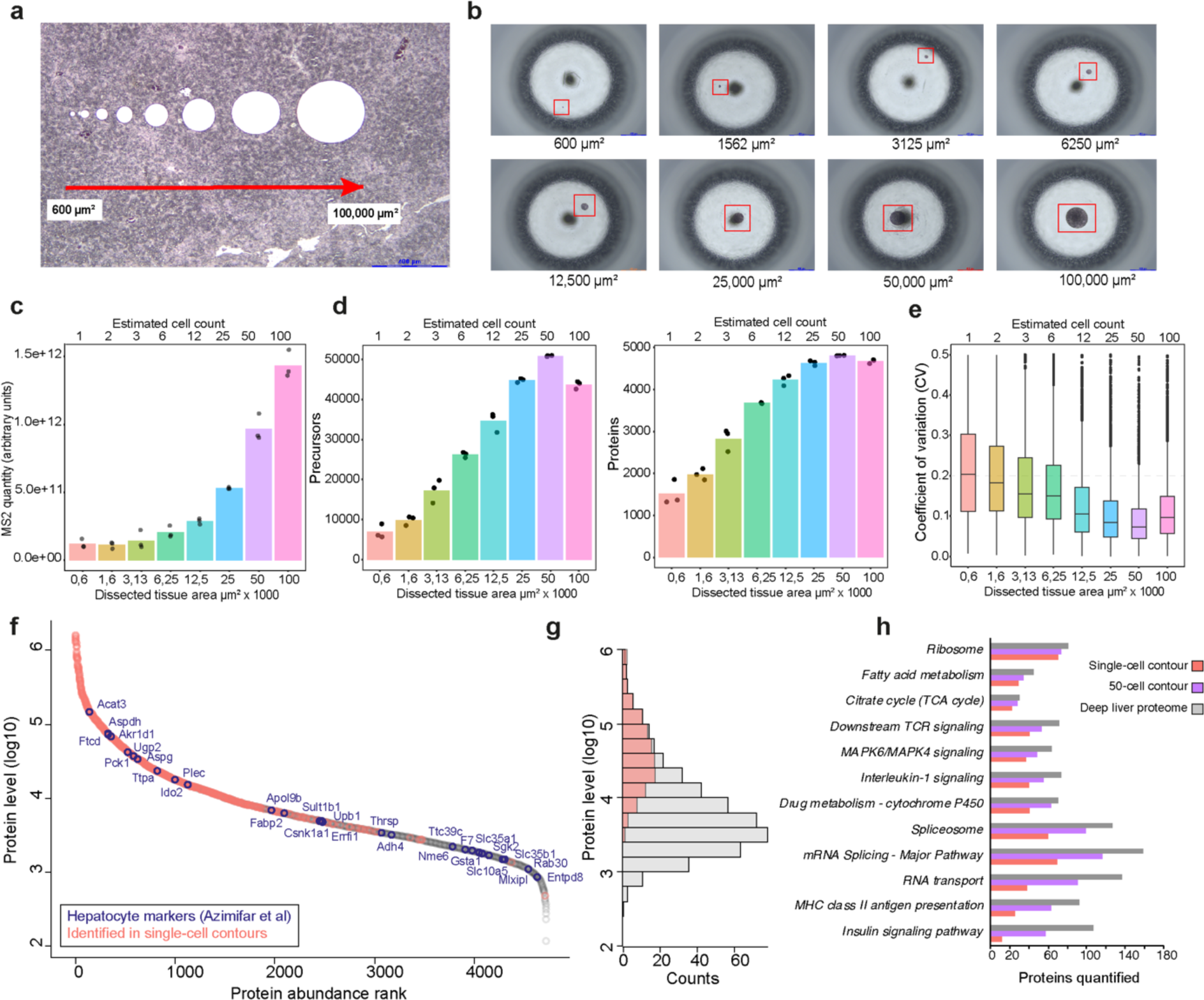
Workflow scalability and single-cell sensitivity. H&E-stained mouse liver tissue section (5 µm). Collected LMD tissue samples are shown from 600 µm^2^ up to 100,000 µm^2^. Single hepatocyte contour isolation (average size 600 µm^2^) was guided by the immunofluorescence signal of the plasma membrane marker Na/K-ATPase and the DAPI signal. Larger regions of interest (ROI) were isolated based on circular contours of pre-defined sizes ranging from 1,500 µm^2^ to 100,00 µm^2^ (2 cell - 100 cells), respectively. Hepatocyte volumes of 4000-6000 µm^3^^33^ were used for cell count estimations of circular ROIs. Scale bar = 400 µm. Tissue inspection after collection into a 384-well low-binding plate. Scale bar = 400 µm. **c-e)** Average MS2 quantities (c), number of quantified precursors (d, left) and proteins (d, right) of the collected tissue samples are shown. For all amounts, three tissue replicates were measured. **e)** Box plots showing the coefficient of variation (CVs) of protein quantification across different tissue areas. CVs were calculated from triplicates of non-log-transformed data. The box plots define the range of the data (whiskers), 25th and 75th percentiles (box), and medians (solid line). **f)** Dynamic range of protein abundance for 50-cell (50,000 µm^2^) contours. Proteins identified in single-cell samples are highlighted, as well as hepatocyte-specific markers ^28^. **g)** Histogram of log10 protein intensities obtained from 600 µm^2^ and 50,000 µm^2^ samples. Proteins identified in single cells cover 2-3 orders of magnitude from the top abundant fraction of the liver proteome. **h)** Reactome and KEGG pathway coverage for single-cell and 50-cell samples. Values show the number of proteins quantified per pathway. For comparison, a deep (> 10,000 proteins) mouse liver dataset ^28^ was included. A minimum of two quantified values from quadruplicate measurements was required for single cells (50% missing values) and three values from triplicates (0% missing values) for the 50-cell samples.

The unprecedented depth of our single FFPE hepatocyte contours, reproducibly quantifying up to 2,000 proteins depending on tissue thickness (**Extended Data Fig. 2a**), encouraged us to further explore these single-cell tissue proteome data. Protein levels generally showed a high degree of concordance with higher loading amounts (**Extended Data Fig. 2f**) and included many known hepatocyte specific markers distributed over a dynamic range of approx. three orders of magnitude (**Fig. 2f-g**). Proteins involved in housekeeping cell functions were quantified at a similar depth of analysis as the 50-cell samples, despite the approx. 33-fold lower total MS2 signal, and, remarkably, compared to the deep mouse liver study covering more than 10,000 proteins ^28^ (**Fig. 2h**). For example, we quantified 70 of 81 ribosomal proteins, 29 of 44 involved in fatty acid metabolism and 22 of 30 TCA cycle-related proteins (**Extended Data Table 2**). Interestingly, while ‘housekeeping’ functions such as ribosomal proteins were more stably expressed, cell metabolism-related proteins showed higher variation in the single-cell contours compared to the 50-cell contours, likely due to proteome averaging (**Extended Data Fig. 2h**). This is in line with a recent liver study revealing strong metabolic differences in single hepatocytes along the liver zonation axis ^18^. For lower abundant pathways, for example, insulin signaling or MHC-2 antigen presentation (12 vs. 107 and 25 vs. 92 detected proteins, respectively) (**Fig. 2h**), a higher discordance in detected proteins was observed between the low and high tissue amounts, making the comprehensive single-cell analysis of these pathways only achievable with further improved workflow sensitivity, or alternatively, by pooling phenotype matched cells, as we conceptualized recently ^15^. We conclude that reproducible single-cell-based FFPE tissue profiling is achievable using optimized sample processing and ultra-sensitive LC-MS workflows revealing important insights into cell identity and function. The scalability of our workflow also enabled the spatially resolved quantification of ∼5,000 proteins from 50-cell regions, capturing a substantial fraction of the cell type specific proteome, thereby complementing single-cell-based analyses.

### Optimizing sample input across tissue and cell types

Based on our tissue dilution experiment in the murine liver, we empirically determined the optimal tissue amount for the highest proteome coverage of small tissue areas using a 15-min active nanoflow gradient combined with dia-PASEF on the Bruker timsTOF SCP. We next addressed how this liver optimum translated to other tissue and cell types and how the recorded MS read-out could be exploited to normalize sample loading from tissue to tissue or cell type to cell type. Such adjustments are of particular importance for ultra-low sample amounts, which are not amenable to peptide concentration measurements routinely used prior to MS bulk analysis. In addition, various factors can affect the total ion current (TIC) derived from different tissue specimens, including sample-related sources of variability such as tissue archival time, which can affect the retrieval of lower abundant proteins ^35^, or biologically due to protein abundance differences across tissue and cell types ^36^. In any case, normalizing and adjusting sample amounts is hence particularly important for low input tissue proteomics and should be carefully assessed. We hypothesized that a simple normalization strategy using the total excised tissue volume (area x section thickness) across specimens should be a poor estimator of the optimal sample amount. To test this, we analyzed a second tissue type, the human FFPE tonsil, which as the secondary lymphoid organ is important for the development of immune tolerance and adaptive immune functions and comprises B and T-cell subsets, as well as other immune and non-immune related cell types ^37^. In the murine liver, 50-100 cell contours (125,000 - 250,000 µm^3^) yielded the highest proteome depth and lowest median protein CVs (**Fig. 2c-e, Extended Data Fig. 3a-c**). Analyzing the same tissue amounts obtained from tonsil, focusing on T and B-cell enriched microregions after immunofluorescence imaging (**Fig. 3a-b**), revealed clearly different sampling optima (**Fig. 3c**), despite a similar total number of quantified proteins per experiment (5,000 – 5,500 proteins, **Extended Data Table 3**). Based on the MS2 signal (sum of MS2 quantities of all peaks) and total quantity (sum of MS2 quantities of identified precursors) retrieved from the DIA-NN output, we estimated that liver on average resulted in roughly three-fold (2.8×) higher MS2 signal compared to tonsil (**Fig. 3d**), likely due to different protein abundances in these two organs (**Fig. 3e**). Importantly, the MS2 signal derived from the liver reference sample was consistent over the tested time frame of six months and independent of section thickness (**Extended Data Fig. 3d**). In other words, a 5-µm thick contour of 25,000 µm^2^ resulted in an almost identical MS2 signal as a 10-µm thick contour of 12,500 µm^2^. The optimal tissue amount for tonsil was 350,000 - 700,000 µm^3^ instead, allowing us to quantify ∼35,000 precursors and ∼5,000 proteins in the 15-min dia-PASEF measurement (**Fig. 3c**), and beyond which no further increase in proteome coverage was apparent. In fact, we observed lower identification rates beyond this saturation point, similar to our findings in liver (**Fig. 2c-e**), further emphasizing the need to carefully assess and normalize sample loading. This prompted us to test if these tissue saturation points obtained from the liver and tonsil data could be exploited to predict the best tissue sampling amount from ‘single-shot’ measurements alone. This simulates a scenario at the beginning of a spatial tissue proteomics study, where a priori proteomics information for a new tissue type is lacking. The accurate prediction of the optimal tissue amount could hence save time and resources and maximize data output. To this end, we laser microdissected replicates of small 60,000 µm^3^ samples (6,000 µm^2^ of a 10 µm slice) of a consecutive tonsil tissue section, such that the obtained intensity was in the linear range of the mass spectrometric signal (>50,000 µm^3^ tissue, **Fig. 3d**) and compared the recorded MS2 precursor quantity to our pre-determined optimum (MS signal = 6-8×10^11^). This way, we extrapolated that 450,000 – 753,000 µm^3^ would be the optimal sampling amount for this tonsil tissue, in excellent agreement with our measured titration data (**Fig. 3c**). We note that this simple normalization strategy is strictly dependent on the employed LC-MS setup and therefore requires empirical data to begin with, but generally recommend to perform such ‘survey’ experiments to pinpoint the optimal tissue amount needed for a specific spatial proteomics research question. In our example, murine liver served as highly consistent reference tissue to predict the ideal tonsil amount, however, other tissue types should be equally suited for such normalization. Moreover, this strategy can prevent TIMS overloading, which can negatively affect proteome coverage and quantification. As a positive byproduct, the acquired raw files of the titration data can also be used to create project-specific refined spectral libraries (Methods), thereby drastically speeding up subsequent DIA-NN searches.

**Figure 3:**
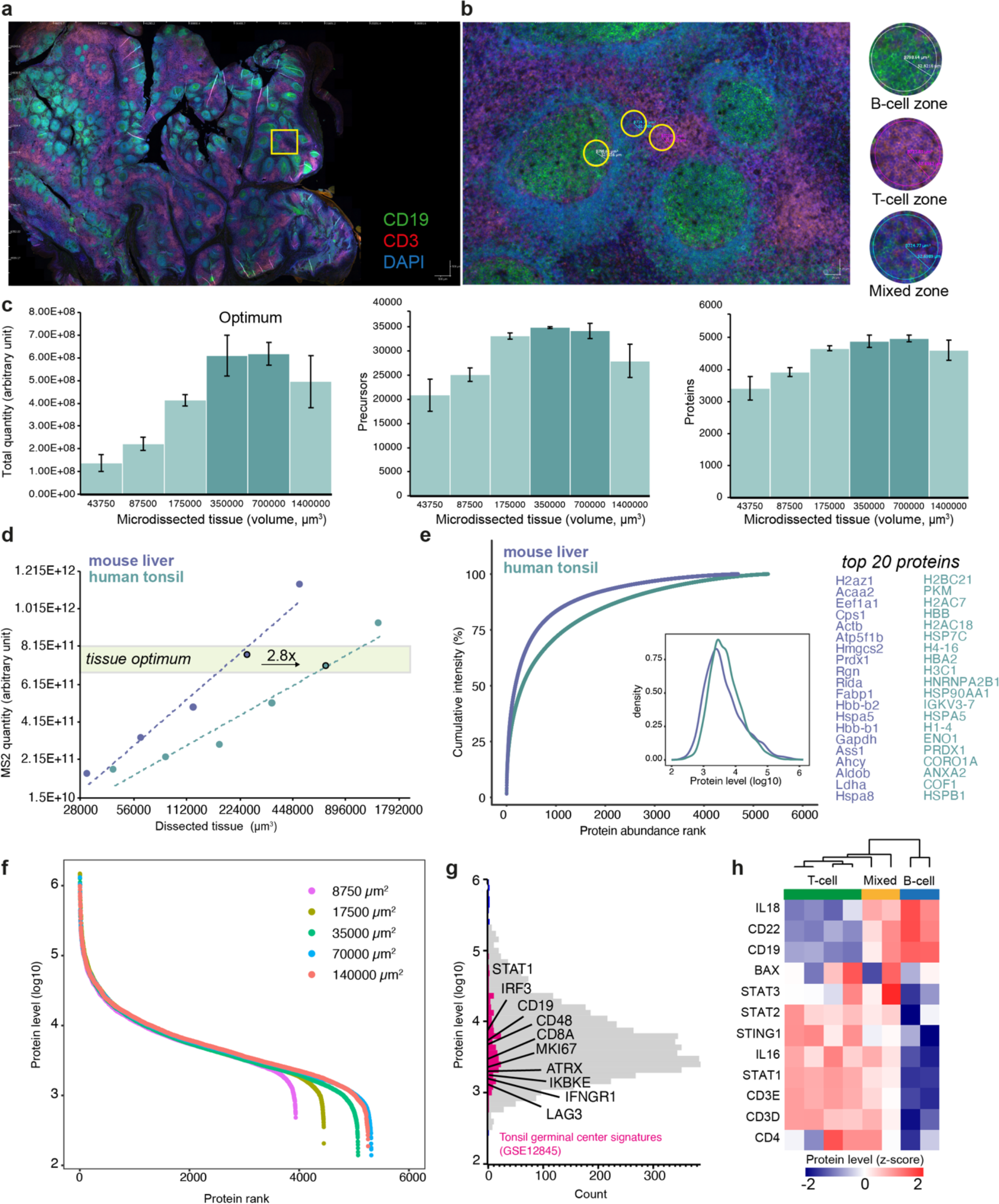
Optimizing sample input across tissue and cell types. Immunofluorescence whole-slide image of a 5-µm thick tonsil tissue section stained for CD3 (T-cells), CD19 (B-cells) and DNA (DAPI). Scale bar = 500 µm. **b)** Magnifications of exemplary B and T-cell enriched regions used for laser microdissection and proteomics profiling. Scale bar = 25 µm. **c)** Bar plots of MS2 intensities (left), precursors (middle) and proteins quantified (right) obtained from increasing amounts of tonsil tissue. Results show averages of six replicates with standard deviations as error bars. Note, the tissue-specific sampling optimum was reached at 350,000 - 700,000 µm^3^, beyond which identifications dropped again. **d)** Comparison of liver and tonsil tissue dilution data. MS2 quantities are plotted against increasing (log2 transformed) tissue amounts (volume in µm^3^). Based on the empirically determined MS2 intensity optimum (MS2 quantity = 6-8E11 for liver and tonsil tissues), 2.8-fold more tonsil tissue was needed to reach the same MS2 quantity. **e)** Cumulative protein intensities of liver and tonsil tissue proteomes ranked from the highest to the lowest abundant protein. Density plot shows protein intensity distributions for both tissues. Top 20 proteins of each tissue are shown on the right. **f)** Dynamic range of protein abundance from different amounts of tonsil tissue. **g)** Histogram of protein intensities of the of 8,750 µm^2^ tonsil tissue sample (43,750 µm^3^ in volume). Proteins highlighted in pink belong to an RNA-seq-based signature of tonsil germinal centers (GSE12845), including known immune and cell type specific markers. **h)** Unsupervised hierarchical clustering of small (6,000 µm^2^ 퀇 10 µm) B-cell, T-cell and mixed B/T-cell regions, related to Fig. 3b. Z-scored protein levels indicate up-regulated (red) or down-regulated (blue) proteins.

As the size of the total dissected tissue area (or a number of collected single-cell contours per sample), which determines the obtained spatial resolution, is inversely correlated with proteome coverage (**Fig. 2c-d**), such survey measurements can also help to find a good balance between these two parameters in the context of the specific research question. To demonstrate this exemplarily, we further analyzed our tonsil proteome data, which in total covered more than 5,000 proteins distributed over four orders of magnitude (**Fig. 3f**). Projecting the different tissue dilution measurements onto the measured dynamic range of protein abundance showed that B or T-cell specific regions of only 8,750 µm^2^ (5-µm thick section) were already sufficient to quantify many key players of immune-cell signaling, cell type specific markers, cytokines and even transcription factors (e.g., STAT1, IRF3, IFNGR1, CD8A, CD19, Ki-67, LAG3, IL-16, IL-18, **Fig. 3b, g-h, Extended Data Fig. 3e-f**) at a spatial resolution of ∼70 - 100 µm (center-to-center, **Fig. 3a**). Consequently, our pipeline should allow the analysis of spatially-resolved proteomes of various B and T-cell niches from regions of as little as ∼4,000 µm^2^ (10-µm thick section).

### Spatially and cell type resolved proteomics of human tonsil tissue

Encouraged by our tonsil titration data, which revealed that known immune cell regulators, cytokines and transcription factors were quantified from tissue regions of as little as 8,750 µm^2^ (43,750 µm^3^ in volume), we systematically investigated the impact of the spatial location on the B and T-cell specific proteomes. Human tonsil represents a prime example of tissues that are organized into distinct microanatomical compartments to fulfill diverse biological functions critical for adaptive immunity. Following antigen encounter of naïve B-cells within the follicle, secondary (activated) follicles are rapidly formed as production centers of antigen-specific B-cells. Following activation, B-cells undergo cycles of maturation and selection within newly formed germinal centers, ultimately giving rise to highly antigen-specific antibody-secreting plasma cells and memory B-cells, the key players of humoral immune response ^38^.

We first mounted a 10-µm thick tonsil section obtained from a patient that underwent bilateral tonsillectomy on a metal frame LMD slide and performed four-color immunofluorescence whole-slide imaging to detect B-cells (CD19), T-cells (CD3), epithelium (pan-CK) and DNA (DAPI) (**Fig. 4a-b**). Using the open-source image analysis software QuPATH ^39^, we selected a total of 146 microregions for automated laser microdissection (Methods) and quantitative proteomics (**Extended Data Table 4**). Based on the well-defined tissue architecture of the human tonsil (**Fig. 4a**), we included small circular regions of 4,000 µm^2^ isolated from primary and secondary B-cell follicles, including subregions of dark, light and grey germinal center B-cell niches, naïve mantel zone derived B-cells, various interfollicular T-cell zones and squamous-cell epithelium (**Fig. 4a-c**). On average, we quantified 1,952 proteins per sample (**Extended Data Fig. 4c**) and 3,334 proteins in total, which resulted in a clear proteome separation by microanatomical region dominated by distinct cell types (**Fig. 4d**). Known cell type markers such as CD3D (T-cell marker), CD19 (B-cell marker) and CDH1 (E-Cadherin, epithelial marker) were highest in the expected sample groups (**Fig. 4e**), confirming high specificity of our proteome data. Germinal centers showed clear signs of increased proliferation (e.g., MKI67, PCNA) and DNA damage (e.g., TOP2A), indicative of active sites of B-cell expansion and somatic hypermutation ^40^. T-cell zones instead featured high levels of the STAT1 transcription factor as well as T-cell modulating cytokines such as IL-16 (**Fig. 4f**). Moreover, the global comparison of all T-cell (n = 34) versus B-cell specific proteomes (mantle zone, n = 36) revealed many bona-fide B and T-cell markers among the top regulated proteins (e.g., CD22, CD3E, CD72, CD5) and potentially many other less characterized ones (**Fig. 4g**). We next focused on different B-cell niches to assess if our spatially resolved data captured known functional differences of spatially defined B-cell zones. Naïve (mantle zone) and activated (germinal center) B-cells showed striking proteome differences (**Fig. 4h**), indicative of their unique biological functions. While mantle zone B-cells were characterized by higher metabolic activity (e.g. TCA cycle and fructose and mannose metabolism), senescence signatures and phosphatidylinositol signaling, activated germinal center B-cells showed strong replication and DNA damage signatures (**Fig. 4h-I, Extended Data Table 4**), in line with their known biological function. Encouraged by this, we next assessed if our quantitative data even separated spatially defined germinal center sub-compartments of dark (sites undergoing active B-cell proliferation and somatic hypermutation) versus light (site of B-cell selection) zones (**Fig. 4j-k**). Clearly, dark zone-derived B-cells featured higher replication and DNA damage response profiles compared to the light zone (**Fig. 4l-m**). The higher expression of the T-cell receptor subunit CD3D in light zones on the contrary illuminated the presence of T-helper cells, important for B-cell selection ^40^. We confirmed this finding by assessing our imaging data, which indeed revealed significantly higher CD3 signals in the light germinal center regions, which was not the case for the B-cell marker CD19 (**Extended Data Fig. 4d**).

**Figure 4:**
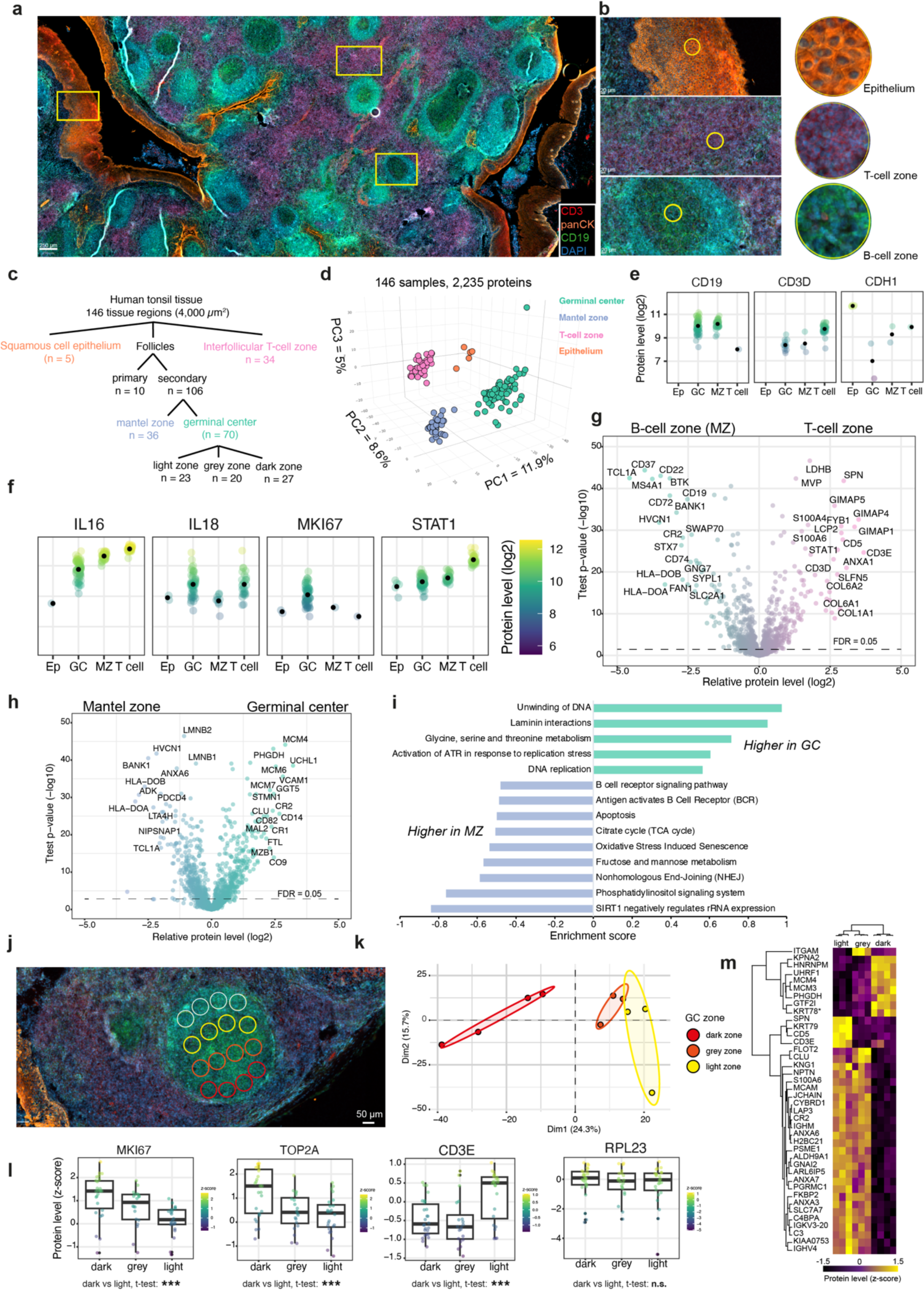
Cell type and spatially resolved proteomics of human tonsil tissue. **a)** Immunofluorescence whole-slide image of a 10-µm thick tonsil tissue section stained for CD3 (T cells), CD19 (B-cells), pan-CK (epithelium) and DNA (DAPI). Scale bar = 250 µm **b)** Magnifications of the exemplary epithelium (EP), germinal center (GC), mantle zone (MZ), and T-cell enriched regions used for laser microdissection and proteomic profiling. **c)** Sample collection strategy and a total number of samples for each tissue region. **d)** 3D PCA of 146 samples based on 2,235 protein groups after data filtering. **e and f)** Log2-protein levels of cell type specific and functional markers quantified in different tissue regions. Black dots indicate average values for each group. **g)** Volcano plot of the pairwise proteomic comparison between the B-cell zone (mantle zone) and T-cell zone. Cell type-specific markers are highlighted in green and turquoise (two-sided t-test, FDR<0.05). **h)** Volcano plot of the pairwise proteomic comparison between mantle zone and germinal center samples. Cell type-specific markers are highlighted in green and blue (two-sided t-test, FDR<0.05). **i)** Pathway enrichment analysis (Reactome and KEGG) based on T-test difference between mantel zone and germinal center samples. Selected pathways with a Benjamin-Hochberg FDR < 0.05 are shown **j)** ROIs used for proteomic profiling of a secondary follicle region. Dark (red), grey (orange) and light (yellow) germinal center zones were selected for proteomic profiling. Mantel zone regions (grey) are shown on top. Scale bar = 50 µm. **k)** PCA of dark, grey and light zone proteomes. Point concentration ellipses are shown for each group with a 95% confidence. **l)** Box-plots of z-scored protein levels for different markers. The box plots define the range of the data (whiskers), 25th and 75th percentiles (box), and medians (solid line). **m)** Unsupervised hierarchical clustering of ANOVA significant proteins (permutation-based FDR < 0.05) from dark, grey and light zone samples. Related to figure 4j and 4k. Protein levels are z-scored showing up-regulated (yellow) and down-regulated proteins (black). *KRT78 is marked as potential contaminant.

In summary, our data delineated how the robust microscopy-guided ultra-low input tissue proteomics workflow introduced here can be applied to study cell type and spatially resolved proteomes in health and disease based on readily accessible archival FFPE specimens.

## DISCUSSION

Spatial tissue proteomics connecting microscopy-based cell phenotyping with laser microdissection guided MS-based proteomics is an emerging discovery concept for the study of cell function and heterogeneity in health and disease. Our group recently co-developed Deep Visual Proteomics, an approach that combines high-parametric imaging and machine-learning-based single-cell phenotyping to guide precise tissue sampling for ultra-sensitive LC-MS analysis. This enabled the profiling of as little as 100 tissue cells per sample to a depth of 3,000 - 5,000 proteins, dependent on the tissue and cell type of interest. However, the flexible and highly modular design of the DVP pipeline, enabling the profiling of single or few cells on one hand, or hundreds of phenotype-matched cells for deeper proteome interrogation (i.e., 5,000 proteins or more) on the other hand, also necessitates carefully designed tissue benchmarking experiments and detailed guidelines to extract most information for diverse biomedical applications. With this in mind, we here introduce, benchmark and apply an optimized ‘end-to-end’ workflow, starting from antigen retrieval comparisons for immunostaining and microscopy, over guidelines for best-practice tissue collection by LMD, towards an optimal nanoflow dia-PASEF LC-MS scheme for highest sensitivity MS-based proteomics. For the first time, we provide evidence that robust ultra-low input proteomics (few or even single excised cells) from FFPE tissue slices is achievable and fully compatible with conventional antigen retrieval and four-marker IF imaging protocols, paving the way for higher-plex IF combinations in the near future. At the same time, our flexible 384-well design makes the protocol easily adaptable to robotic workflows for further protocol automation. We find that low microliter volumes (∼2 µl) for sample preparation are sufficient to achieve true single-cell sensitivity, which makes this workflow principally accessible to any laboratory. On the Bruker timsTOF SCP, this allowed us to quantify up to 2,000 proteins from single hepatocyte contours and around 5,000 proteins from 50-cell regions, demonstrating excellent workflow scalability. On the timsTOF Pro2 instrument, this translated to nearly 1,000 high-confidence proteins from 1-2 hepatocytes (∼1,500 µm^2^ regions) and a somewhat similar proteome depth for the 50-cell samples, further emphasizing the broad applicability of our protocol. The data from single excised hepatocytes also revealed that a ∼25-µm spatial resolution is principally achievable for tissue types such as liver, on par with the spatial resolution of state-of-the-art spatial transcriptomics ^41, 42^. These data also show that highly abundant ‘housekeeping’ proteins, such as ribosomal proteins, are quantifiable at a depth and precision similar to much higher sampling amounts, for example from large-scale bulk measurements. Our data also show that many metabolic pathways (e.g., TCA cycle or fatty acid metabolism) are likewise amenable to single-cell-based proteomic analysis, as also shown recently ^18^. Notably, ‘housekeeping’ related proteins are often poorly correlated to mRNA abundances^10, 43^, making these pathways particularly attractive for early applications of single-cell-based tissue proteomics.

We also provide guidance on how to empirically determine the optimal tissue amount to achieve the highest proteome coverage, while avoiding instrument overloading, which is particularly pertinent for single-cell sensitivity MS setups. We introduce a simple normalization strategy using the extracted MS2 signal directly from the search results to find the optimal tissue amount for literally any low-input spatial tissue proteomics experiment. This strategy can also be applied to normalize for qualitative sample differences, which are often observed for archival specimens due to, for example, varying archival times ^30^. The importance of this can be illustrated through the eyes of a pathologist. For the histomolecular analysis of cancer progression states, samples are typically distributed over several FFPE tissue blocks (for example pre-cancer, primary tumor and metastasis), potentially collected over many years. Survey experiments, as outlined here using the example of human tonsil tissue, could guide the best sampling strategy for deep and reproducible LMD-assisted proteomics.

However, while the latest generation MS instruments feature excellent sensitivity when combined with optimized sample preparation workflows, such as the one introduced here, MS throughput is still a major bottleneck for tissue sections that are often larger than 1 cm^2^. Isobaric or non-isobaric multiplexing strategies generally offer good alternatives to label-free based methods further increasing sample throughput ^44–46^, but they are still not sufficient to deal with this tremendous analytical bottleneck. One can estimate that for the profiling of one 1cm 퀇 1cm tissue section, sampled by non-overlapping 50 µm 퀇 50 µm squares, amounts, which we here show in liver and tonsil tissues to be amenable to the reproducible quantification of 2,000 - 3,000 proteins, 40,000 measurements would be required for gridded whole-slide sampling schemes. This is far beyond the reach of current LC-MS setups, which typically analyze 20-50 tissue proteomes per day. Instead, the integration of whole-slide IF imaging for detailed cell and cellular neighborhood phenotyping offers a powerful strategy for the prioritization of cells and ROIs subjected to global proteome analysis. We illustrate this in tonsil tissue, where we use four-marker whole-slide IF imaging to guide the sampling of over 140 microregions per single batch, covering naïve and activated B-cell niches, interfollicular T-cell zones and squamous cell epithelium. From only ∼63 µm 퀇 63 µm regions (4,000 µm^2^ regions), we quantified spatially resolved proteomes of activated germinal center niches, illuminating sites of clonal B-cell expansion and somatic hypermutation. Intriguingly, based on these 1,500 – 3,000 protein measurements per microregion, we quantified key players of immune cell function including cytokines and transcriptional regulators, emphasizing the power of our ‘biological’ fractionation strategy to dig deep into the cell type and spatially-resolved tissue proteome.

In conclusion, we here provide a detailed and optimized framework for MS-based spatial tissue proteomics of ultra-low-input archival specimens combining high-content imaging, laser microdissection and ultra-sensitive mass spectrometry.

## ACKNOWLEDGEMENTS

We would like to thank our colleagues at the Max Delbrück Center (MDC) for their support and fruitful discussions. In particular, we thank Ulrike Stein for her support to perform the mouse liver experiments. Philipp Mertins and Simon Haas we thank for their critical feedback on the manuscript and Christian Sommer for mass spectrometry support. We also thank Andreas Mund (Center for Protein Research, University of Copenhagen) and Lisa Schweizer (MPI of Biochemistry, Munich) for their input and feedback on the LMD membrane comparisons. We thank Simon Schallenberg (Charité Pathology, Berlin) for his help with the tonsil tissue experiments. Furthermore, we acknowledge the MDC technology platforms ‘Proteomics’ and ‘Advanced Light Microscopy’ for their great support. All authors acknowledge support by the Federal Ministry of Education and Research (BMBF), as part of the National Research Initiatives for Mass Spectrometry in Systems Medicine, under grant agreement no. 161L0222.

## AUTHOR CONTRIBUTIONS

Conceptualization: A.M and F.C.; Methodology: A.M., D.Q., J.K., J.N., F.C.; Experiments: A.M., D.Q., S.F., J.K., F.C.; Data analysis: A.M. and F.C.; Figures: A.M. and F.C.; Supervision: F.C; Funding acquisition: F.C; Writing the original draft: F.C. All the authors reviewed and edited the manuscript.

## COMPETING INTEREST STATEMENT

The authors declare no competing interests.

## METHODS

### Mouse experiments and organ harvesting

For the mouse liver proteome experiments, 6-8 weeks old female C57BL/6 mice from Jackson Laboratory were used. C57BL/6 mice were housed in individually ventilated cages in a specific pathogen-free mouse facility at the Max-Delbrück Center for Molecular Medicine (Berlin, Germany).

For liver excision, anesthetized mice were sacrificed by cervical dislocation, and the livers were removed, rinsed twice in ice-cold PBS, and transferred to 4% formaldehyde solution for fixation (fixation for at least 24h to 48h). Thereafter, livers were paraffin-embedded for further histological analyses. The animal experiments were performed in accordance with the United Kingdom Coordinated Committee on Cancer Research (UKCCR) guidelines and were approved by local governmental authorities (Landesamt für Gesundheit und Soziales Berlin, Germany).

### Human tissue samples

Tonsil tissues were obtained from two female patients aged 34 and 36. Both presented with recurrent chronic tonsillitis and underwent bilateral tonsillectomy. The pathological examination revealed hypertrophy and hyperplasia of the lymphoid follicles as well as enlarged germinal centers. Furthermore, an abundance of collagen fibers within the stroma could be observed. There were no signs of active inflammation or malignancy.

Resected tonsils were fixed in 10% buffered formalin before gross processing. After overnight fixation, the specimens were weighed, measured, and macroscopically evaluated. Afterwards, the specimens were cut into 5-µm-thick slices and two representative slices were embedded in paraffin for further microscopic diagnosis. After histological examination the tissue blocks were stored at room temperature at the archive of the Institute of Pathology at Charité University Hospital, Campus Mitte. The study was performed according to the ethical principles for medical research of the Declaration of Helsinki and approval was obtained from the Ethics Committee of the Charité University Medical Department in Berlin (EA1/222/21).

### Hematoxylin-eosin staining

Briefly, PEN glass slides (Carl Zeiss, 15350731) were treated with UV light for 1 hour. PPS frame slides (Leica, 11600294) were used directly for the next steps. FFPE tissue sections were cut with a microtome (5 µm or 10 µm-thick), air dried at 37 °C overnight and heated at 60°C for 10 minutes to facilitate better tissue adhesion. Next, tissue sections were deparaffinized by washing 2×5 minutes in Neo Clear (Sigma Aldrich, 1.09483.5000), followed by a series of 99%, 80% and 70% ethanol for 2 minutes, respectively, and rehydrated by immersing in milliQ water three times. Then, slides were stained in Mayer’s hematoxylin for three minutes and immersed in tap water for another ten minutes, rinsed in milliQ water and stained with eosin for 30 seconds. Subsequently, the slides were dehydrated by submerging in 70%, 80%, and 99% ethanol serially. Samples were finally air-dried and stored at RT until imaging. Before imaging, a cover glass (Corning, CLS2980223, #1.5) was mounted with Aqua Poly Mount medium (Polysciences Europe GmbH, 18606-20).

### Immunofluorescence staining

Following tissue sectioning, mounting on LMD slides and de-paraffinization (described above), two epitope retrieval methods were compared: heat-induced (HIER) and protease (pepsin)-induced (PIER, 10 min) epitope retrieval. HIER was done by submerging in EnVision FLEX Target Retrieval Solution High pH solution (diluted to 1X) (Agilent Dako, cat.no. K8004) and heating in a steamer at 95°C for 20 min, and subsequently cooled down in a pre-heated PBS buffer at room temperature for 30 min. Odyssey Blocking Buffer (LI-COR BioScience, 927-70001) was used for blocking in a humidified chamber for 30 min at room temperature. PIER was done by applying pepsin (Agilent Dako, cat.no. S3002) on the tissue slide for 5 minutes at 37°C, and immersed in PBS solution to stop the reaction. The slides were washed two times with PBS and air dried before antibody staining.

For murine liver tissue stains, the primary antibody targeting Na/K-ATPase (dilution 1:100, Abcam, ab76020) was diluted in Odyssey Blocking Buffer and incubated overnight at 4 °C in a humidified chamber. Next, tissue specimens were washed 3x in PBS and secondary antibodies for the visualization of Na/K-ATPase (Alexa Fluor 488 donkey anti-rabbit, dilution 1:250, A32790, Invitrogen) were diluted in Odyssey Blocking Buffer and applied for 1 hour at room temperature in the dark. After staining, slides were washed 3x in PBS, counterstained by Hoechst (dilution 1:1000 in PBS, Thermo Fisher Scientific, #62249) for 10 minutes and followed by three washes in PBS and two washes in milliQ. Before imaging, a cover glass (#1.5) was mounted using ProLong^TM^ Diamond anti-fade mounting medium (Thermo Fisher Scientific, P36961).

For tonsil tissue stains following tissue sectioning, mounting on LMD slides and de-paraffinization, the tissues were subjected to heat-induced epitope retrieval as described above. Odyssey Blocking Buffer was used for blocking in a humidified chamber for 30 min at room temperature. Next, conjugated primary antibodies targeting CD20 (dilution 1:50, Thermofisher, 53-0202-80, Alexa Fluor 488), CD3 (dilution 1:100, Abcam, ab198937, Alexa Fluor 647), and pan-cytokeratin (dilution 1:100, Thermofisher, 41-9003-80, eFluor 570) were diluted in Odyssey Blocking Buffer (and incubated overnight at 4 °C in a humidified chamber. Tissue specimens were washed 4x in PBS, counterstained by Hoechst (dilution 1:1000 in PBS, Thermo Fisher Scientific, 62249) for 10 minutes, washed 4x in PBS and 2x in milliQ water. Subsequently, the slides were dehydrated by submerging them in 70%, 80%, and 99% ethanol serially. Samples were finally air-dried and stored at RT until imaging. Before imaging, a cover glass was mounted with ProLong^TM^ Diamond anti-fade mounting medium.

### High-resolution microscopy

Images of immunofluorescence-labeled tonsil tissue sections were acquired using an Axioscan 7 system (Zeiss), equipped with wide-field optics, a Plan-A photochromat 10x/0.45 M27 objective and a quadruple-band filter set for Alexa fluorescent dyes. The wide-field acquisition was performed using the Colibri 7 LED light source and an AxioCam 712m camera. Images were obtained automatically with Zeiss ZEN 3.7 (blue edition) at non-saturating conditions (16-bit dynamic range).

Images of immunofluorescently-labeled and hematoxylin-eosin stained murine liver tissue sections were acquired on the Leica LMD7 system using the LAS X software (version 3.7.524914, Leica Microsystems), an HC PL FLUOTAR 10x/0.32 DRY objective and the DF7000T camera. The microscope was equipped with the following filter sets: LMD-Dapi, LMD-Cy3, LMD - Alexa594, LMD-YFP, LMD-CY5, LMD-BGR, GFP.

### Image analysis and contour export for laser microdissection

Image analysis was performed in QuPath (version 0.4.3) and BIAS (BioStudies Archive accession number S-BSST820). Annotations of different regions of interest were manually created in QuPath (version 0.4.3) after image analysis. Three clearly recognizable reference points (x-y pixel coordinates) were also assigned needed for precise contour transfer between the screening and laser microdissection microscopes. Contours were then exported as geojson file and formatted into an xml file compatible with the Leica LMD7 software. The processing was performed in a jupyter notebook using geopandas (Version 0.12.2). A detailed open-source workflow for precise contour transfer is under Github release.

### Laser microdissection

We used the Leica LMD 7 system and Leica Laser Microdissection V 8.3.0.08259 software for the collection of tissue contours. Depending on the contour size, tissue was cut with a 20x or 63x objective in fluorescence or brightfield mode. The following laser settings were used for the 20x objective (HC PL FL L 20x/0.40 CORR): power 56, aperture 1, speed 15, middle pulse count 1, final pulse -1, head current 37 – 45% (depending on tissue type and section thickness), pulse frequency 801 and offset 101. For the 63x objective (HC PL FLUOTAR L 63x/0.70 CORR XT), the settings were: power 59, aperture 1, speed 50, middle pulse count 1, final pulse -1, head current 45-50%, pulse frequency 3000, offset 101.

Contours were cut and sorted into a low-binding 384-well plate (Eppendorf 0030129547) configured over the ‘universal holder’ function with one empty well between samples.

### Sample preparation for LC-MS analysis

A detailed sample preparation protocol is provided in the **Supplementary information**. Tissue samples were collected by manual cutting or by automated cutting after contour import into low-binding 384-well plates. For mouse liver tissue samples (5- or 10-μm-thick sections cut with a microtome), regions of 600 μm^2^ - 100,000 μm^2^ were collected. To concentrate tissue pieces at the bottom of each well after LMD collection, 15 µl of acetonitrile was added to each well, briefly vortexed and vacuum dried (15min at 60°C). Another well inspection is recommended before proteomics sample preparation to ensure high collection efficiency.

We tested three different protocols, DDM-based, ACN-based and a combination of DDM and ACN. The lysis buffer for the DDM-based protocol consisted of 0.1% DDM, 5mM TCEP, 20mM CAA and 0.1M TEAB in water. 2µl of lysis buffer was added to each sample well using the MANTIS Liquid Dispenser (Formulatrix, V3.3 ACC RFID, software version 4.7.5) and the high-volume diaphragm chips (Formulatrix, cat.no. 233128). The plate was closed with a PCR ComfortLid (Hamilton), and heated at 95°C for 60 minutes. Then, samples were shortly cooled down, and 1µl of LysC was added (prediluted in water to 2 ng/µl) and digested for minimum 2 hours at 37°C in the thermal cycler (50°C lid temperature). Subsequently, 1µl of trypsin was added (prediluted in water to 2 ng/µl) and incubated overnight at 37 °C in the thermal cycler. The next day, digestion was stopped by adding trifluoroacetic acid (TFA, final concentration 1% v/v), and samples were vacuum dried before peptide clean-up.

For the ACN-based protocol, the lysis buffer consisted of 5mM TCEP, 20mM CAA, 0.1M TEAB diluted in water. 2µl of lysis buffer was added to each sample well, the plate was closed with PCR ComfortLid, and heated at 95°C for 60 minutes in a thermal cycler (Bio-Rad S1000 with 384-well reaction module) at a constant lid temperature of 110°C. Next, 1ul of 100% ACN was added to each well, and the plate was incubated for an additional 60 min in a thermal cycler at 75°C at a constant lid temperature of 110°C. Then, samples were shortly cooled down, and 1µl of LysC was added (prediluted with 0.1M TEAB, 30% ACN water to 2 ng/µl) and digested for 2 hours at 37°C in the thermal cycler (50°C lid temperature). Subsequently, 1µl of trypsin was added (prediluted with 0.1M TEAB, 10% ACN water to 2 ng/µl) and incubated overnight at 37 °C in the thermal cycler. The next day, digestion was stopped by adding trifluoroacetic acid (TFA, final concentration 1% v/v), and samples were vacuum dried before peptide clean-up.

For the combined DDM/ACN-based protocol, the lysis buffer for the DDM-based protocol consisted of 0.1% DDM, 5mM TCEP, 20mM CAA and 0.1M TEAB in water. 2µl of lysis buffer was added to each sample well using the MANTIS Liquid Dispenser and the high-volume diaphragm chips. The plate was closed with a PCR ComfortLid, and heated at 95°C for 60 minutes. Then, samples were shortly cooled down, and 1µl of LysC was added (2 ng/µl in 0.1M TEAB (pH 8.5) and 30% ACN in milliQ water) and digested for minimum 2 hours at 37°C in the thermal cycler (50°C lid temperature). Subsequently, 1µl of trypsin was added (2 ng/µl containing 10% ACN and 0.1M TEAB (pH 8.5) in milliQ water.) and incubated overnight at 37 °C in the thermal cycler. The next day, digestion was stopped by adding trifluoroacetic acid (TFA, final concentration 1% v/v), and samples were vacuum dried before peptide clean-up.

### Peptide clean-up with C-18 tips

Evotip (Evosep, Odense, Denmark) based peptide clean-up was performed as recommended by the manufacturer. Briefly, 20 ul of buffer B (99.9% ACN, 0.1% FA) was added to each C-18 tip (EV2013, Evotip Pure, Evosep) and centrifuged at 700 rpm for 1 minute. Then, 20ul of buffer A (99.9% water, 0.1% FA) was added from the top of each C-18 tip, activated in isopropanol for 20 seconds and centrifuged again at 700 rpm for 1 minute. Digested tissue samples were then loaded onto Evotips, washed once with 20ul buffer A and finally eluted with 20ul buffer B to a 96-well plate (Thermo Fisher Scientific, AB1300), and vacuum dried (15min at 60°C). Samples were stored at −20 °C until liquid chromatography–mass spectrometry (LC–MS) analysis. For LC-MS analysis, 4.2 µl of MS loading buffer (3% acetonitrile, 0.1% TFA in water) was added, the plate was vortexed for 10 seconds and centrifuged for 1 minute at 700g. 4 µl were finally injected into the mass spectrometer.

### Liquid chromatography–mass spectrometry (LC – MS) analysis

LC-MS analysis was performed with an EASYnLC-1200 system (Thermo Fisher Scientific) connected to a trapped ion mobility spectrometry quadruple time-of-flight mass spectrometer (timsTOF SCP and timsTOF Pro2, Bruker Daltonik) with a nano-electrospray ion source (CaptiveSpray, Bruker Daltonik). The autosampler was configured to pick samples from 384- and 96-well plates.

Peptides were loaded on a 20-cm home-packed HPLC column (75-µm inner diameter packed with 1.9-µm ReproSil-Pur C18-AQ silica beads, Dr. Maisch).

Peptides were separated using a linear gradient from 7-30% buffer B (0.1% formic acid and 90% ACN in LC-MS grade water) in 14 minutes, followed by an increase to 60% for 1 minute and a 1.5-minute wash in 90% buffer B at 250 nl min^-1^. Buffer A consisted of 0.1% formic acid in LC-MS grade water. The total gradient length was 21 minutes. A column oven was used to keep the column temperature constant at 40°C.

For dia-PASEF analysis, we used a dia-PASEF method with 8 dia-PASEF scans separated into 3 ion mobility windows per scan covering a 400-1000 m/z range by 25 Th windows and an ion mobility range from 0.64 to 1.37 Vs cm^-2^. The mass spectrometer was operated in high sensitivity mode, with an accumulation and ramp time at 100 ms, capillary voltage set to 1750V and the collision energy as a linear ramp from 20 eV at 1/K0 = 0.6 Vs cm^-2^ to 59 eV at 1/K0 = 1.6 Vs cm^-2^. The collision energy was ramped linearly as a function of ion mobility from 59 eV at 1/K0 = 1.6 V s cm^-2^ to 20 eV at 1/K0 = 0.6 V s cm^-2^.

### Proteomics raw data analysis

We used DIA-NN^27^ (version 1.8.1) for dia-PASEF raw file analysis and spectral library generation.

#### Spectral library generation

For spectral library generation to analyze dia-PASEF data, human and mouse FASTA files were downloaded from Uniprot (2022 release, UP000000589_10090 Mus Musculus, UP000005640_9606, downloaded on April 10th and April 8th 2022 respectively). DIA-NN in silico predicted libraries were generated by providing the human or mouse FASTA file and frequently found contaminants ^47^ (mouse + mouse tissue contaminants, or human + universal contaminants). Deep learning-based spectra, RTs and IMs prediction were enabled for the appropriate mass range of 300-1200 m/z. N-terminal M excision was enabled and cysteine carbamidomethylation was enabled as a fixed modification. A maximum of 2 missed cleavages was allowed, and the precursor charge set to 2 -4. For the generation of project-specific refined libraries, in-silico-generated mouse and human libraries were used to search 20-50 raw files of high-sample amounts, such that the optimal total ion current was reached (see Fig. 2 and 3). The refined murine liver library consisted of 68,006 precursors, 61,554 elution groups and 8,225 protein groups. The refined human tonsil library consisted of 47,999 precursors, 44,675 elution groups and 8,137 protein groups.

#### Search of dia-PASEF raw files with refined libraries

DIA-NN was operated in the default mode with minor adjustments. Briefly, MS1 and MS2 accuracies were set to 15.0, scan windows to 0 (assignment by DIA-NN), isotopologues were enabled, no MBR, heuristic protein inference and no shared spectra. Proteins were inferred from genes, neural network classifier was set to single-pass mode, quantification strategy as ‘Robust LC (high precision)’. Cross-run normalization was set to ‘RT-dependent’, library generation as ‘smart profiling’, speed and Ram usage as ‘optimal results’.

### Statistical analysis

Proteomics data analysis was performed with Perseus ^48^ (version 1.6.15.0) and within the R environment (https://www.r-project.org/, version 4.2.2) with the following packages: ggplot2 (v3.4.2), FactoMineR (v2.8), factoextra (v 1.0.7.999), plotly (v4.10.1), reshape2 (1.4.4), viridis (v0.6.3). For differential expression analysis (t-test or ANOVA, Figs. 3 and 4), data were filtered to keep only proteins with 70% non-missing data in at least one group. Missing values were imputed based on a normal distribution (width = 0.3, downshift = 1.8) before statistical testing. For multi-sample (ANOVA) or pairwise proteomic comparisons (two-sided unpaired t-test), a permutation-based FDR of 5% was applied to correct for multiple hypothesis testing. Principal component analysis was performed in R (see packages above). 1D pathway enrichment analysis ^49^ (Fig. 4i) was done in Perseus based on the KEGG (https://www.genome.jp/kegg/) and Reactome Pathway Database (reactome.org), using a Benjamini-Hochberg FDR cut-off of 0.05. The minimum category size was set to 5. Venn diagrams were created using online tool (https://bioinformatics.psb.ugent.be/webtools/Venn/).

### Data availability

The mass spectrometry proteomics data have been deposited to the ProteomeXchange Consortium (http://proteomecentral.proteomexchange.org) via the PRIDE partner ^50^ with the dataset identifier PXD042367. Raw files will be accessible upon publication.

## EXTENDED DATA

**Extended Data Figure 1, related to Fig. 1.**
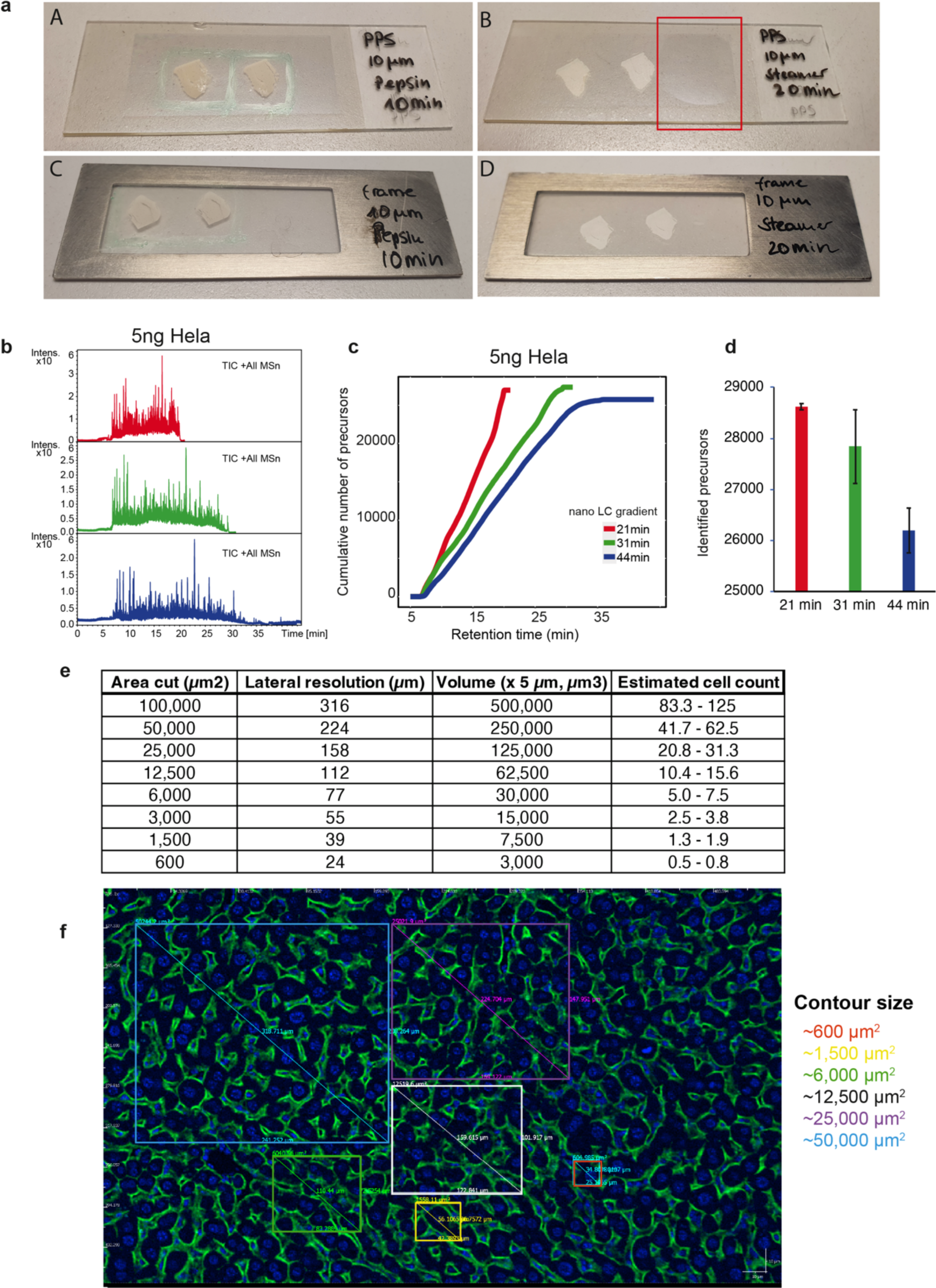
**a)** PPS glass membrane and PPS metal frame slides after heat-induced (HIER) or protease-induced (PIER) epitope retrieval. The red box shows an area of membrane distortion after HIER. **b)** Total ion chromatogram (TIC) of three chromatographic gradients (21min, 31min, 44min) obtained from 5ng HeLa (Pierce) dia-PASEF runs. **c)** Cumulative number of identified precursors from the 21min, 31min and 44min runs as a function of retention time in minutes. **d)** Number of identified precursors of the three chromatographic gradients. **e)** Summary of the tissue dilution experiment of mouse liver FFPE tissue. Hepatocyte sizes of 4000-6000 µm^3^^33^ were used for cell count estimations. **f)** Immunofluorescence image of mouse liver tissue stained against the plasma membrane marker Na/K ATPase and DAPI (DNA). Boxes exemplarily show different sizes of regions used for laser microdissection and proteomic profiling. Scale bar = 10 µm.

**Extended Data Figure 2, related to Fig. 2.**
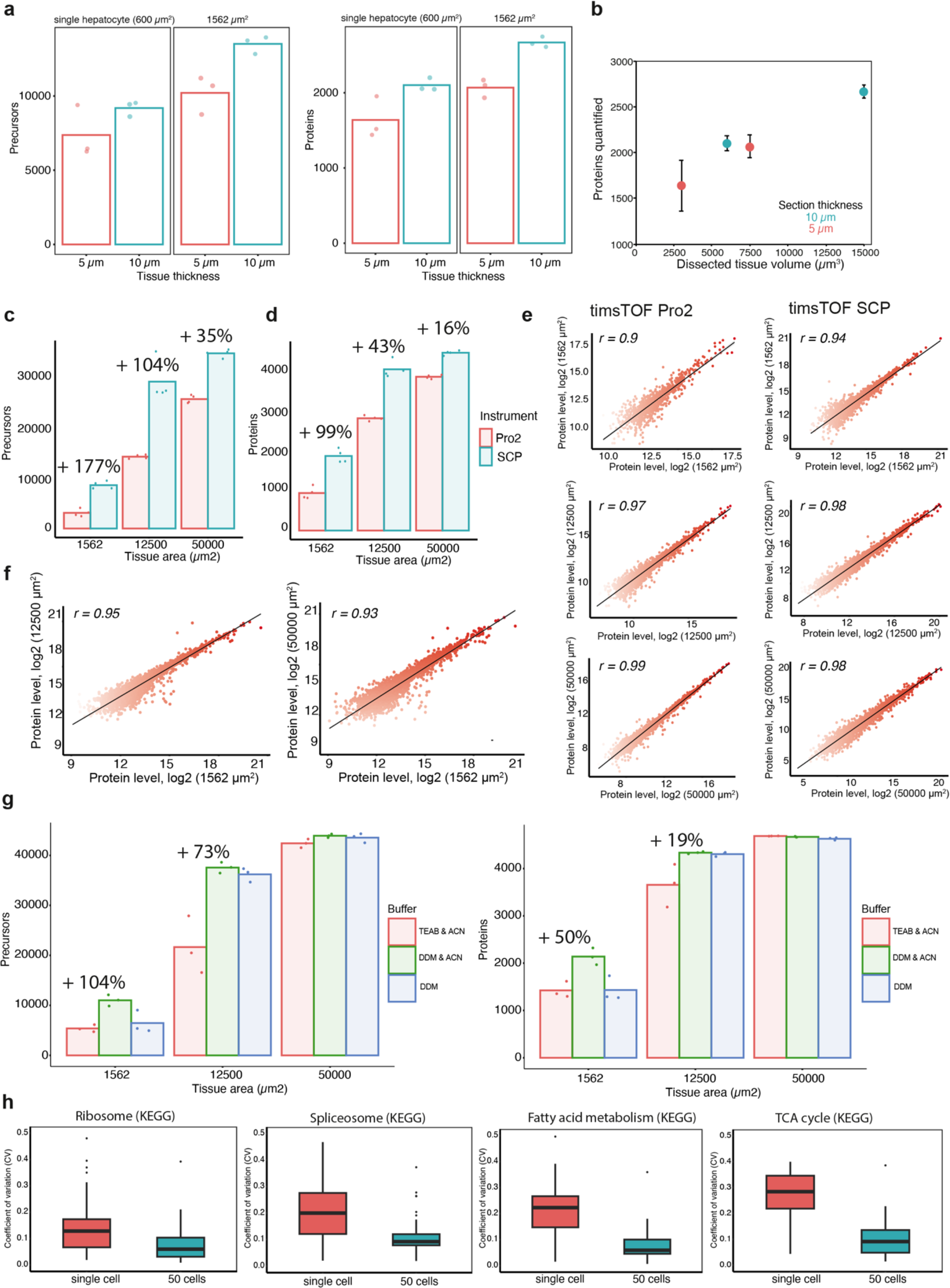
**a)** Average precursor and protein identifications from utralow-input murine liver tissue samples of 5 µm and 10 µm thickness. **b)** Average number of identified proteins as a function of dissected tissue volume. **c) and d)** Comparison of timsTOF Pro2 and timsTOF SCP data. Precursor (c) and protein (d) identifications from murine liver tissue samples. Areas of 1500 µm^2^, 12500 µm^2^ and 50000 µm^2^ of a 5-um thick section were compared. **e)** Proteome correlations of different tissue areas measured in dia-PASEF mode on the timsTOF Pro2 and timsTOF SCP instruments. Pearson correlations and p-values are shown. **f)** Proteome correlations between different tissue amounts (1500 µm^2^, 12500 µm^2^ and 50000 µm^2^) obtained on the timsTOF SCP instrument. Pearson correlations and p-values are shown. **g)** Impact of different lysis buffers on proteome coverage. Areas of 1500 µm^2^, 12500 µm^2^ and 50000 µm^2^ of a 5-um thick mouse liver section were profiled using three different methods based on organic solvent only (ACN), DDM only (no ACN) or combined (DDM & ACN) protocol. **h)** Box plots showing coefficient of variations (CVs) of protein quantifications from four different KEGG pathways. CVs were calculated from non-logarithmic data of quadruplicate measurements for 1500 µm^2^ and 50000 µm^2^ regions. The box plots define the range of the data (whiskers), 25th and 75th percentiles (box), and medians (solid line). Outliers are plotted as individual dots outside the whiskers. The bar plots in panels a, b, c, d and g show average values from minimum triplicate measurements.

**Extended Data Figure 3, related to Fig. 3.**
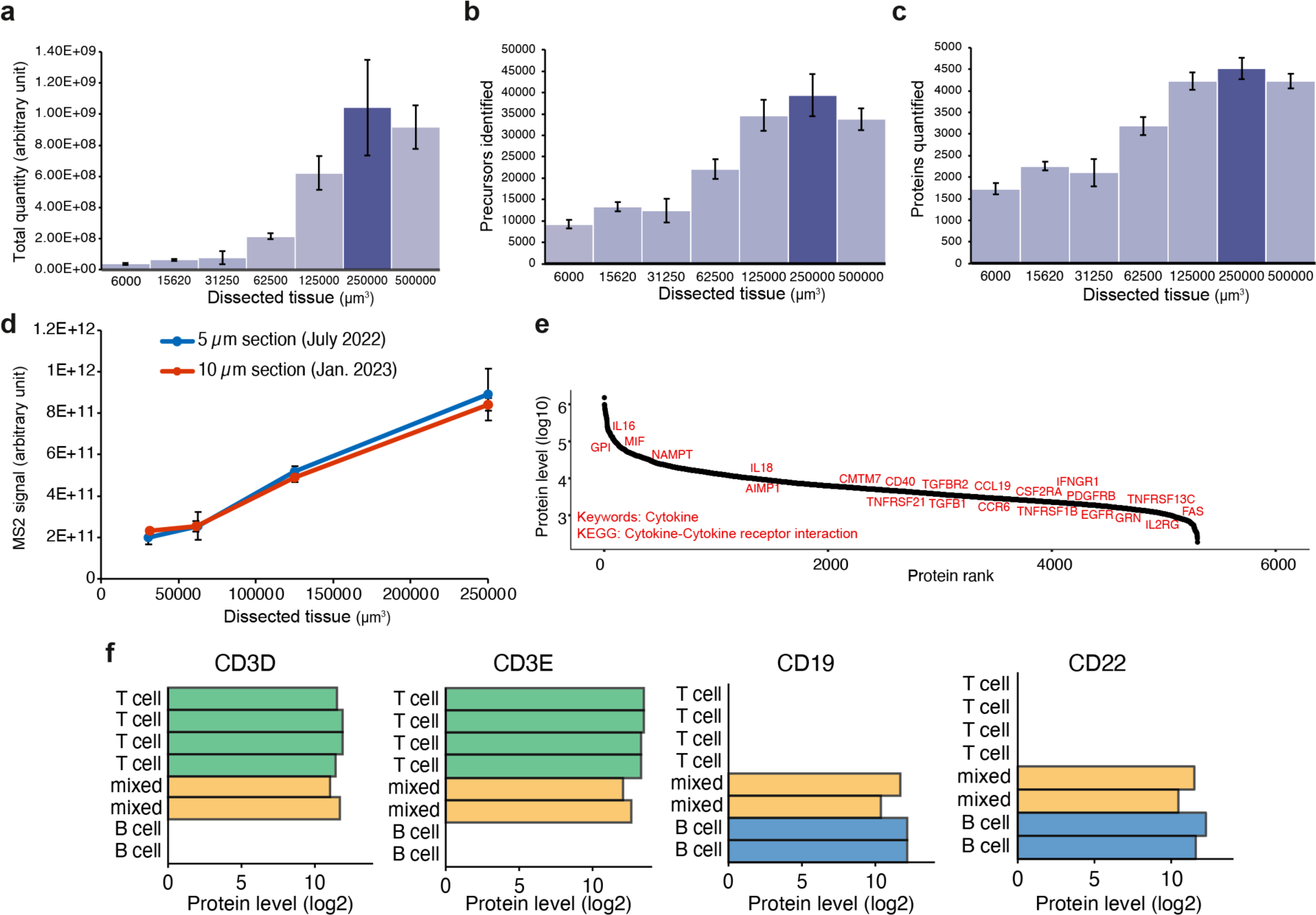
**a), b) and c)** Bar plots showing total quantities (sum of precursor MS2 intensities) (a), identified precursors (b), and identified proteins (c) from increasing amounts of murine liver tissue. Results show averages of quadruplicate measurements with standard deviations as error bars. Note, the tissue specific sampling optimum was reached at 250,000 µm^3^, beyond which identifications dropped. **d)** MS2 signals obtained from the liver reference tissue show consistent results over the measured time frame of six months. Note, nearly identical signals were measured for volume (µm^3^) matched tissue samples of 5 µm and 10 µm thickness. **e)** Dynamic range of protein abundance (log10) for tonsil tissue. Proteins were ranked by abundance level. Cytokines (Keywords) and Cytokine-Cytokine receptor interaction (KEGG) related proteins are highlighted. **f)** Bar plots showing relative protein levels (log2) of T-cell (CD3D, CD3E) and B-cell (CD19, CD22) markers obtained from T-cell, B-cell and mixed zones of human tonsil tissue. Related to Fig. 3b and 3h.

**Extended Data Figure 4, related to Fig. 4.**
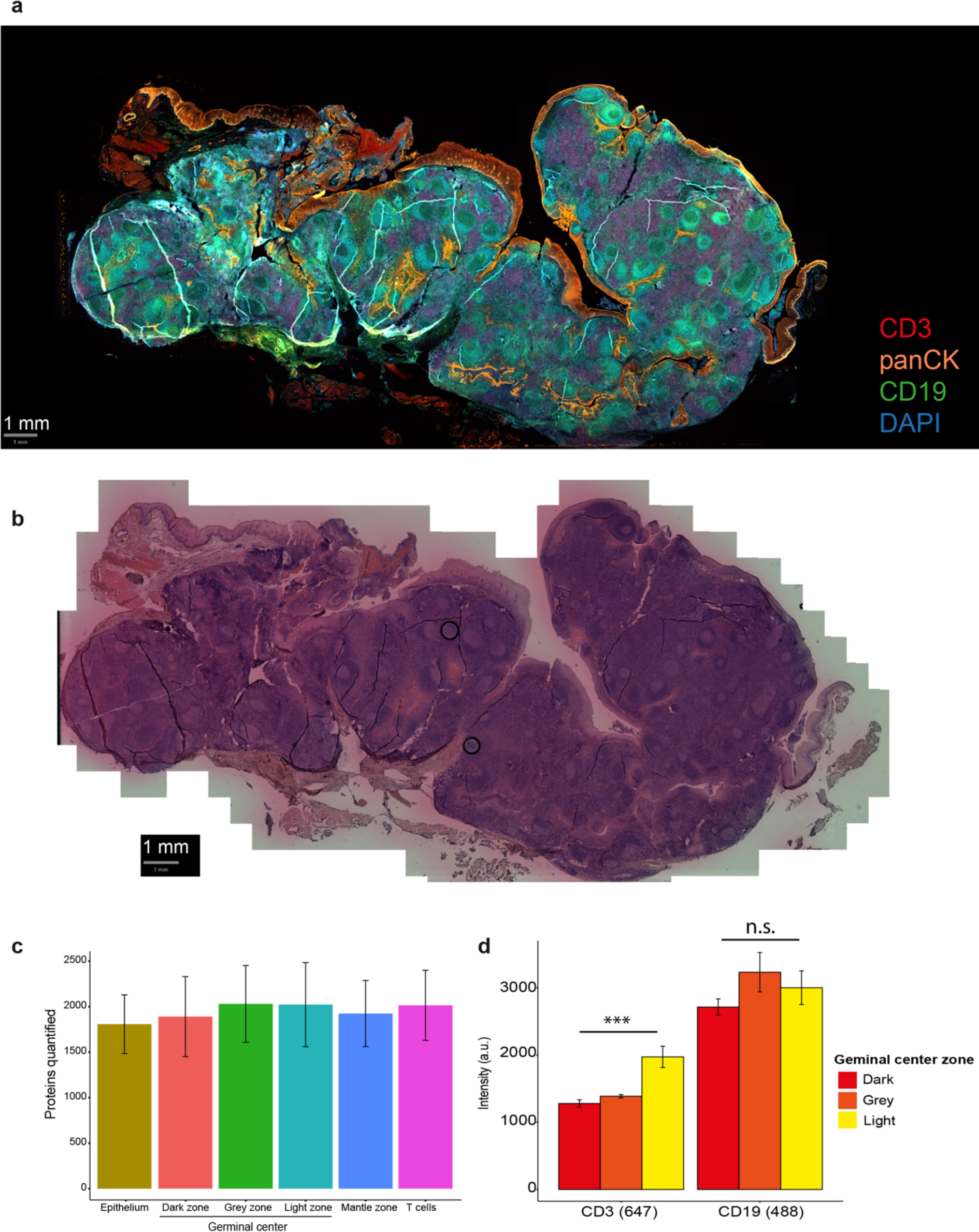
**a)** Immunofluorescence whole-slide image of a 10-µm thick tonsil tissue section stained for CD3 (T cells), CD19 (B-cells), pan-CK (epithelium) and DNA (DAPI). Scale bar = 1 mm. **b)** Brightfield image of the same tonsil tissue section (panel a) after H&E staining. Scale bar = 1 mm. **c)** Number of protein quantifications obtained from different tonsil microregions of 4,000 µm^2^. **d)** Intensities derived from CD3 (647) and CD19 (488) channels from contours in light, grey and dark germinal center zones depicted in Fig. 4j. CD3 intensities showed significant differences between dark and light zone samples (two-sided T-test, p-value < 0.01), but no significance for CD19 intensities (p-value > 0.05).

## Notes

### Competing Interest Statement

The authors have declared no competing interest.

### Summary of Updates

The legends of Figs. 2c-e and 2h were corrected.

