## Supplementary information for "A framework for ultra-low input spatial tissue proteomics"

**<u>Disclaimer:</u> This protocol includes hazardous chemicals and any use of it is strictly at your own risk (!). Please read safety information.**

### Detailed sample preparation protocol

#### REAGENTS

• Triethylammonium bicarbonate pH 8.5 (TEAB; Merck, cat.no. T7408-100ML)

• (!) n-Dodecyl-beta-Maltoside (DDM)

• (!) Acetonitrile (ACN; HPLC-grade, VWR, cat.no. 83640.290)

• (!) Trifluoroacetic acid (TFA; Sigma-Aldrich, cat.no. 96924-250ML-F)

• Endoproteinase Lys-C (Promega, cat.no VA1170)

• Proteomics grade modified trypsin (Promega, cat.no V5117)

• (!) Isopropanol (ISO, Sigma-Aldrich, cat.no. 1070222511)

• (!) Tris(2-carboxyethyl)phosphine hydrochloride (TCEP; Sigma-Aldrich, cat.no. C4706-2G)

• (!) 2-chloroacetamide (CAA; Sigma-Aldrich, cat.no. C4706-2G )

• (!) Formic acid (FA; Merck Millipore, cat.no 1.00264.1000)

• (!) EnVision FLEX Target Retrieval Solution High pH (50X) (Agilent Dako, cat.no. K8004)

• Super PAP-pen liquid blocker mini (Science Services, cat.no. N71312-N)

• Microscopy Neo-Clear (xylene substitute) (Sigma Aldrich, cat.no. 1.09843.5000)

• Phosphate buffer saline PBS for IF staining

• Odyssey Blocking Buffer (LI-COR BioScience, cat.no. 927-70001)

• Prolong Diamond antifade mounting medium (Invitrogen, cat.no. P36961)

• Aqua Poly Mount (Polysciences Europe GmbH, cat.no. 18606-20)

• Pepsin solution for antigen retrieval (Agilent Dako, cat.no. S3002)

• Cover glass (Corning, cat.no. CLS2980223, #1.5)

• PPS frame slides (Leica, cat.no. 11600294)

• PEN glass slides (Carl Zeiss, cat.no 15350731)

#### EQUIPMENT

• Thermo Scientific Microm HM 355S (Thermo Fisher Scientific,)

• Eppendorf ThermoMixer (Bio-Rad S1000 with 96- and 384-well reaction module)

• 384 well low-binding plate (Eppendorf, cat.no. 0030129547)

• 96 well plate (Thermo Fisher Scientific, cat.no. AB1300)

• Evotip Pure (Evosep, cat.no. EV2013)

- Microtome blade N35 (VWR, cat.no. 720-2369)
- Evaporative concentrator: Eppendorf Vacuum Concentrator Plus with 96-well plate rotor
- MANTIS Liquid Dispenser (Formulatrix, V3.3 ACC RFID, software version 4.7.5)
- High-volume diaphragm chips for MANTIS Liquid Dispenser (Formulatrix, cat.no. 233128)
- PCR ComfortLid (Hamilton)
- Steamer (any steamer that can reach 100°C can be used)

### REAGENT SETUP

- **Stock solutions:**

0.1 μg/μl trypsin protease (dilute in milliQ water)

0.2 μg/μl LysC protease (dilute in milliQ water)

550 mM 2-chloroacetamide (CAA) in milliQ water

500 mM tris(2-carboxyethyl)phosphine hydrochloride (TCEP) in milliQ water

5% DDM in milliQ water

These buffers can be aliquoted at -20 °C.

*CAUTION: CAA and TCEP are toxic. Pipette them in a fume hood and handle with gloves.*

- **Lysis buffer**

Prepare 0.1% DDM buffer containing 5mM TCEP, 20mM CAA, 100mM TEAB pH 8.5 in milliQ water

*IMPORTANT: This buffer should be freshly prepared.*

*CAUTION: CAA and TCEP are toxic. Pipette them in a fume hood and handle with gloves.*

- **Digestion buffer**

LysC: Prepare 2 ng/μl in 0.1M TEAB and 30% ACN in milliQ water.

Trypsin: Prepare 2 ng/μl containing 10% ACN and 0.1M TEAB (pH 8.5) in milliQ water.

*IMPORTANT: These enzyme dilutions should be freshly prepared.*

*CAUTION: ACN is toxic. Pipette it in a fume hood and handle with gloves.*

• **MS loading buffer**

3% ACN, 0.1% FA (vol/vol) in milliQ water. This buffer is stable at RT.

*CAUTION: FA acid vapor is a severe irritant of eyes, mucous membranes, and skin. When preparing this buffer, pipette under fume hood and wear gloves.*

**Tissue cutting using a microtome**

1. FFPE tissue blocks are stored at room temperature.
2. Place the tissue block into the microtome holder.
3. Attach clean microtome blade.
4. Cut tissue sections 5-10 μm thick.
5. Mount sections on PPS or PEN membrane slides by slowly lifting up the membrane side of the slide from underneath the floating tissue section. For metal slides, use the flat side of the slide.
6. Dry the slides at 37 °C overnight.

Important guidelines to consider when cutting formalin-fixed paraffin-embedded (FFPE) tissue blocks are the following:

• Only use demineralized or distilled water to prevent calcification of water bath.

• Microtome blades are sharp. To minimize the risk of being injured, always place the knife guard over the microtome blade.

• The flotation bath was filled with ultrapure water and set to 50°C. Warm water is critical to allow tissue to smooth out before being mounted on a frame slide.

• When cutting tissue, periodically clean the microtome blade from excess paraffin that can affect tissue cutting.

• For increased tissue adherence, PEN glass slides should be treated with UV light for 1 hour.

#### Tissue deparaffinization and hematoxylin-eosin (H&E) staining

Prior to tissue collection by laser microdissection (LMD), FFPE samples were deparaffinized and immunofluorescence (following epitope retrieval) or H&E stained as described below. Note, H&E staining can also be performed after the final round of immunofluorescence staining (see Supplementary Fig. S4a-b).

| **Deparaffinization** | | | **H&E staining** | |
|---|---|---|---|---|
| **Step** | **Time** | | **Step** | **Time** |
| Oven at 60 °C | 10 min | | Mayer's Hematoxylin | 3 min |
| Neo-Clear | 10 min | | Water | 10 min |
| 2 x 99 % ethanol | 2 min each | | 2 x milliQ | short dip |
| 80 % ethanol | 2 min | | Eosin | 30 seconds |
| 70 % ethanol | 2 min | | 2 x milliQ | short dip |
| 2x milliQ | 2 min each | | 70 % ethanol | 10 dips |
| Water | 10 min | | 80 % ethanol | 10 dips |
| | | | 99 % ethanol | 10 dips |
| | | | 99 % ethanol | 10 dips |

#### Antigen retrieval for immunofluorescent staining

**a) Heat-induced**

1. After deparaffinization and rehydration, transfer tissue slides to a plastic staining jar filled with EnVision FLEX Target Retrieval Solution High pH (diluted to 1x) (pH 9).
2. Heat in the steamer for 20 min at 95 °C.
3. Let cool down in pre-warmed PBS buffer.
4. Wash the slides by immersing in PBS buffer.
5. Air dry at room temperature and continue with immunofluorescence staining.

**b) Pepsin-induced**

1. Thaw pepsin reagent at 37°C.
2. Pipette 500 $\mu$l of pepsin reagent directly onto the tissue slide and incubate for 10 min at 37°C.
3. Wash slides by immersing in PBS buffer.
4. Air dry at room temperature and continue with immunofluorescence staining.

**Tissue immunofluorescence staining (murine liver example)**

1. After tissue cutting, mounting on LMD slides, de-paraffinization and antigen retrieval, the slides are air-dried at room temperature.

   *NOTE: Outlining tissue sections with a PAP-pen is helpful to prevent spreading of antibody solution over the frame slide.*

2. Wash the slides by immersing in PBS buffer.
3. Block the slides in Odyssey Blocking Buffer in a moist chamber. Apply a 500 μl droplet, incubate 30 min at room temperature.
4. Apply primary antibody targeting Na/K-ATPase (dilution 1:100 in Odyssey blocking buffer, Abcam, ab76020). Incubate at 4°C overnight.
5. Next day, wash the tissue slides by immersing in PBS buffer.
6. Apply secondary antibodies for the visualization of Na/K-ATPase (Alexa Fluor 488 donkey anti-rabbit, dilution 1:250 in Odyssey blocking buffer, A32790, Invitrogen). Incubate at room temperature for 1 hour.

   *IMPORTANT: Antibody solutions should be prepared freshly.*

7. Wash the tissue slides by immersing in PBS buffer.
8. Counter stain with Hoechst (dilution 1:1000 in PBS) for 10 min.
9. Wash the tissue slides by immersing in PBS buffer.
10. Wash the tissue slides by immersing in milliQ water.
11. Perform dehydration steps by immersing in 70%, 80%, 99%, 99% EtOH each 1 min.
12. Mount coverslip using hydrophilic mounting medium and perform imaging.

    *IMPORTANT: After imaging, it is recommended to remove the coverslip as soon as possible, as long attachment will result in difficulties removing it later and may affect tissue quality. For immunofluorescence, we recommend ProLong Diamond antifade hydrophilic mounting medium.*

13. After imaging, remove coverslip through rotation in PBS in a staining jar. After removal, dip slide in water and shortly air-dry. Store samples at 4°C until laser microdissection.

For laser microdissection, the collection plate (low-binding 384-well plate) should always be covered by a lid (e.g. Hamilton ComfortLid or Nunc Aluminium Seal Tape, Thermo Scientific, #232699), unless it is inside the LMD system, to avoid contamination. Collector inspection should be done to ensure that cut tissue pieces are at the bottom of the well plate. Small contours (e.g. single cells) can stick to the side wall of the well. Here, it is helpful to add 15-20 $\mu$l of ACN to the wells after collection and immediately centrifuge in an evaporative concentrator. This will concentrate the collected material at the bottom of the well to allow low volume sample preparation.

*NOTE: It is advantageous to leave one well empty between samples of a 384-well plate. This way all samples per row can be easily processed with a multichannel pipette.*

#### Sample preparation for proteomics

Pipetting steps can either be performed manual (e.g. multi-channel pipette) or using robotic workflows such as the Formulatrix Mantis system as described below.

#### Buffer dispensing with MANTIS

1. Turn on MANTIS Liquid Dispenser and start software. Make sure that the pressure values are in the acceptable range.
2. Install the the high-volume diaphragm chip to the chip port.
3. Run wash of the chip by clicking "Wash Input" button. The washing can be done with water or with 70% ethanol.
4. Load the appropriate dispense list.
5. Load the buffer into 1ml pipette tip and load the tip to the chip.

*IMPORTANT: Avoid spilling of the buffer on top of the chip. Liquid can enter the MANTIS Air Ribbon and solenoid bank resulting in damaging of the instrument.*

6. Put the destination plate onto the plate holder (384-well plate), select the target wells and input the dispense volume in µl.
7. Select the chip holding the pipette with buffer by clicking the corresponding chip.
8. Start the run to dispense.

   *IMPORTANT: It was observed that the first few dispensing are not precise; therefore, it is recommended to first dispense a few µl into empty wells before dispensing into the wells containing samples.*
9. After dispensing, carefully remove the pipette tip from the chip.
10. Run wash of the chip by clicking "Wash Input" button. The washing can be done with water or with 70% ethanol.
11. For storage, manually pipette a few µl of 30% Glycerol into the chip.

***1. Tissue lysis, protein reduction, alkylation and tryptic digestion overnight (day 1, ~16 h)***

☐ Add 2 µl lysis buffer and close with ComfortLid. Incubate for 1 hour at 95°C in a thermal cycler at a constant lid temperature of 110°C.

*CRITICAL: Make sure that the plate is tightly closed. Evaporation of the lysis buffer may result in incomplete tissue lysis and affect subsequent steps.*

☐ Add 1 µl of 2 ng/µl lys-C to each well containing tissue sample. Centrifuge shortly to make sure that the enzyme is in the buffer. Incubate at 37°C in the thermal cycler for minimum 2 hours at a constant lid temperature of 50°C.

*CRITICAL: When adding enzyme to the well, one can pipette it to the side wall and centrifugation will make sure that enzyme will reach the sample without disturbing the tissue pieces at the bottom of the well.*

☐ Add 1 μl of 2 ng/μl trypsin to each well containing tissue sample. Centrifuge down to make sure that the enzyme is in the buffer. Incubate at 37°C in the thermal cycler overnight at a constant lid temperature of 50°C.

*NOTE: Occasionally, buffer evaporation may occur, which results in slightly different sample volumes after heating. If necessary, adjust sample volumes with milliQ water.*

***2. Peptide clean-up (day 2, ~1 h)***

☐ After overnight digestion, add TFA to a 1% final concentration to acidify the solution and inactivate trypsin and LysC. Mix by vortexing for 10 seconds.

*NOTE: At this stage the plate can be frozen or continue with the next steps.*

☐ ***Peptide purification via Evotips (follow manufacturer's protocol)***

- Prepare 3 solvent containers filled with buffer A (0.1% FA in water), buffer B (0.1% FA in ACN) and isopropanol.
- Pipette 100 μl of isopropanol into the wells of a clean 96-well plate.
- Place Evotips into the Evotip box corresponding to the positions of where the samples will be eluted
- Transfer 20 μl of buffer B to Evotips. Place the Evotips in the centrifuge with appropriate counterbalance and centrifuge 700 g for 60 seconds.

  *CRITICAL: Empty box of buffer B after centrifugation steps.*
- Place the Evotip adaptor rack on top of the 96 well plate filled with isopropanol. Make sure that the tips are immersed in isopropanol and soak for a minimum of 10 seconds.

*CRITICAL: Make sure that all tips are pale white after soaking. If the tips are not pale, soak for additional 10-20 seconds.*

- Place the Evotips still in the adapter rack to the empty Evotip box. Transfer 20 µl of buffer A to each Evotip. Place the Evotips in the centrifuge with appropriate counterbalance and centrifuge 700 g for 60 seconds.

  *CRITICAL: From now on it is very important to keep the Evotips wet. During loading stage, always keep them in the chamber filled with buffer A, make sure that the tips are immersed.*

- Transfer 20 µl of the sample to each Evotip.

  *NOTE: To maximize sample transfer efficiency, it is recommended to load samples twice by adding 10 µl buffer A each for both transfer steps.*

- Wash each Evotip by adding 20 µl of buffer A. Place the Evotips in the centrifuge with appropriate counterbalance and centrifuge 700 g for 60 seconds.

  *NOTE: if an Evosep system is used for LC-MS analysis, stop here and fill up tips with enough buffer A to prevent drying out. Strictly follow recommendations from the Evosep manual. If plate-based LC-systems are used (e.g. Easy-nLC 1200, Thermo Fisher Scientific), continue with peptide elution into 96-well plates.*

- Transfer Evotips in the adapter rack to a clean 96-well plate. Add 20 µl of buffer B to each Evotip. Place the Evotips in the centrifuge with appropriate counterbalance and centrifuge 700 g for 60 seconds.
- Dry samples in the evaporative concentrator. After drying the samples can then be dissolved in 2-4 µl MS loading buffer and analyzed.
